# USP39 promotes hepatocellular carcinogenesis through regulating alternative splicing in cooperation with SRSF6/HNRNPC

**DOI:** 10.1101/2023.01.24.525324

**Authors:** Jingyi Zheng, Shasha Wu, Mao Tang, Shaoyan Xi, Yanchen Wang, Jun Ren, Hao Luo, Pengchao Hu, Liangzhan Sun, Yuyang Du, Hui Yang, Fenfen Wang, Han Gao, Ziwei Dai, Xijun Ou, Yan Li

## Abstract

Abnormal alternative splicing (AS) caused by alterations in spliceosomal factors is implicated in cancers. Standard models posit that splice site selection is mainly determined by early spliceosomal U1, U2 snRNPs. Whether and how other mid/late-acting spliceosome components such as USP39 modulate tumorigenic splice site choice remain largely elusive. We observed that hepatocyte-specific knock-in of USP39 promoted hepatocarcinogenesis and potently regulated splice site selection. In human liver cancer cells, USP39 promoted tumor proliferation in a spliceosome-dependent manner. USP39 depletion deregulated hundreds of AS events, including the oncogenic splice-switching of KANK2. Mechanistically, we developed a novel RBP-motif enrichment analysis and found that USP39 modulated exon inclusion/exclusion by interacting with SRSF6/HNRNPC in both humans and mice. Our data represent a paradigm for the control of splice site selection by mid/late-acting spliceosome proteins and their interacting RBPs. USP39 and possibly other mid/late-acting spliceosome proteins may represent potential prognostic biomarkers and targets for cancer therapy.

**Graphic Abstract:** 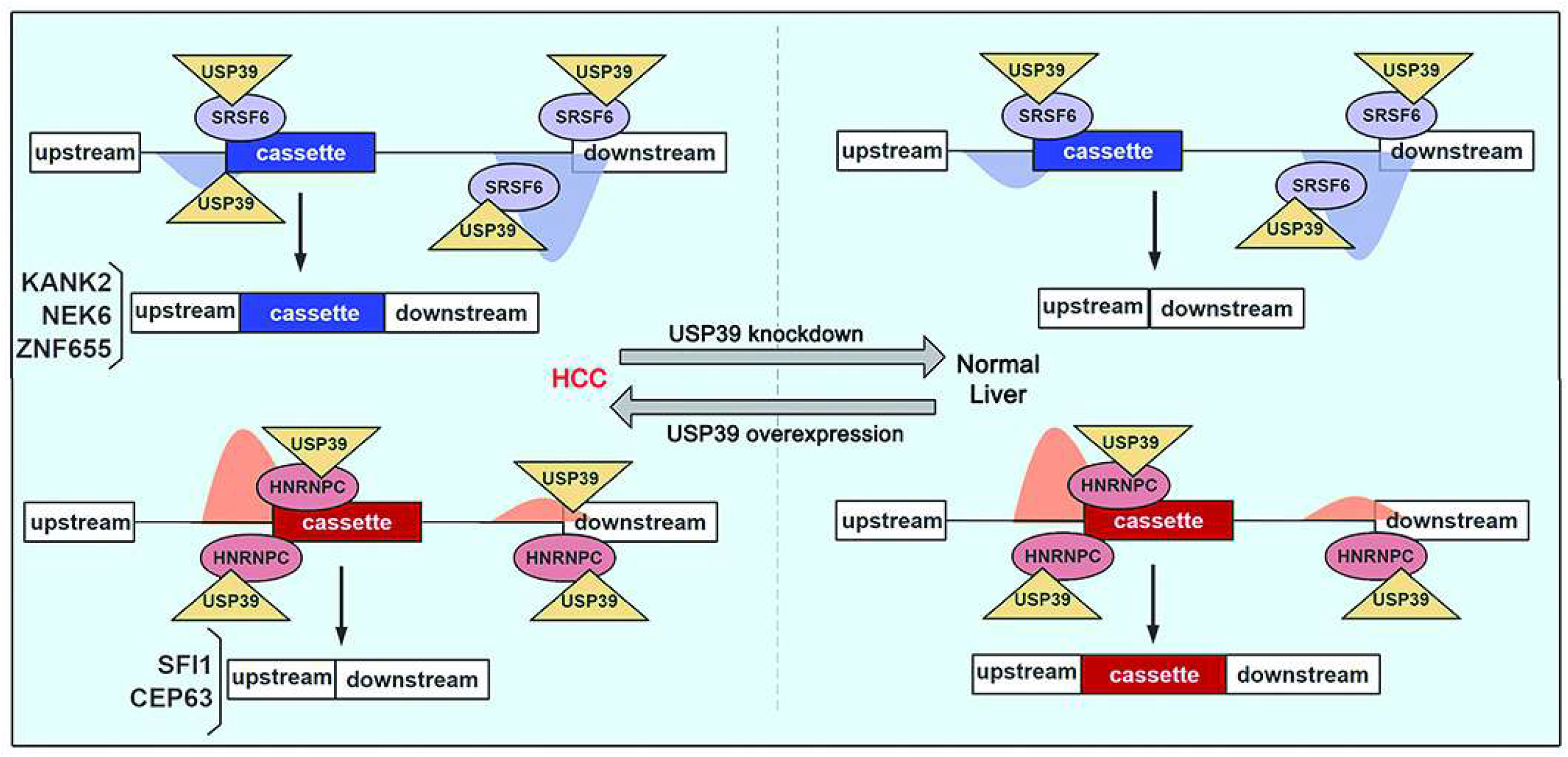

## Introduction

Alternative splicing (AS) is an important mechanism of RNA processing that generates different mRNA/protein isoforms from a single mRNA transcript, thereby enlarging the diversity and complexity of the transcriptome and proteome. Pre-mRNA splicing is carried out by the spliceosome, a megadalton complex comprising five small nuclear ribonucleoprotein particles (U1, U2, U4/5/6 snRNP) and over 100 core set proteins. The key of AS regulation lies in the selection of splicing sites, which is believed to be mainly determined by U1, U2 snRNPs and their interacting RNA binding proteins (RBPs). Current models posit that U1 snRNP recognizes the 5’ splice site, and U2 snRNP auxiliary factor (U2AF) recognizes sequences at the 3’ end of introns. U2AF binding helps to recruit U2 snRNP to the upstream branch site, forming complex A. Additional interactions that regulate splice site selection are mediated by RBPs (e.g., SR and hnRNP protein families), which recognize auxiliary sequences in the pre-mRNA to promote or inhibit complex A assembly. Subsequent binding of preassembled U4/5/6 tri-snRNP forms complex B, which undergoes a series of conformational changes to form complexes Bact and C, and concomitantly carries out the two trans- esterification reactions to generate splicing intermediates and products. (Will & Lührmann, 2011)

Over the past decade, rapid developments in high-throughput technologies have revealed broad alterations in splicing in various cancers.(H. Chen et al., 2019; Seiler, Peng, et al., 2018; Yu et al., 2020) Genetic alteration and/or abnormal expression of spliceosomal components have been frequently detected and contribute to the abnormal splicing patterns in tumors. These findings indicate that cancer-related isoforms and various splicing regulatory factors can be used as potential targets in cancer therapy, leading to a new treatment strategy called spliceosome-targeted therapies (STTs).(J. Chen & Weiss, 2015; Danan-Gotthold et al., 2015; Eymin, 2021) U1 and U2 snRNP components have been intensively studied and pharmacologically targeted because of their direct influence on splicing site recognition and frequent mutation in hematological tumors.(Alsafadi et al., 2016; Seiler, Yoshimi, et al., 2018) Small molecules targeting these components, such as SF3B inhibitor, cause severe toxic side effects due to its general regulation of splicing efficiency, although they are effective in various cancers.(Eskens et al., 2013) On the other hand, targeting non-spliceosomal regulators such as RBM39 is less toxic, however its efficacy in cancer patients is limited. The current dilemma prompted researchers to turn to other mid- or late-acting spliceosome components for solution. This may be especially true for cancer types like hepatocellular carcinoma (HCC), because extensive abnormal expression of spliceosomal components rather than mutation of U1/U2 snRNP components is frequently detected in these cancers.

Several key issues need to be addressed whether mid- and late-acting spliceosome components can be used as therapeutic targets. Despite extensive deregulation of spliceosomal gene expression, *in vivo* models and data are needed to determine whether these proteins are drivers or mere passengers in cancer pathogenesis. Considering standard models of AS regulation at early steps of spliceosome assembly, whether and how mid- and late-acting spliceosome components modulate splice site choice, rather than only affecting efficiency of splicing. Take Ubiquitin-specific protease 39 (USP39), a component of U4/U6. U5 tri-snRNP, as an example. USP39 is an ortholog of yeast Sad1 and was first identified to facilitate U4/U6.U5 tri-snRNP assembly and complex B formation and conversion to Bact.(Bertram et al., 2017; Lygerou, Christophides, & Séraphin, 1999) It is also defined as a member of the deubiquitylation family, but is considered to be devoid of deubiquitinating activity due to the absence of three crucial catalytic residues. The aberrant expression of USP39 has been implicated in the tumorigenesis of various malignancies.(Fraile et al., 2017; Zhu et al., 2022) Overexpression of USP39 facilitates constitutive splicing of individual pre-mRNAs in different cancers.(Ding et al., 2019; Y. Huang et al., 2016; S. Wang et al., 2021; Xiao et al., 2022; Zhao et al., 2021) In addition, USP39 regulates SP1 and ZEB1 protein stability via deubiquitylation.(X. Dong et al., 2021; Li et al., 2021) However, as a mid/late-acting spliceosome protein, the roles and targets of USP39 in alternative splicing regulation remains unclear. And little is known about the mechanism underlying USP39-mediated splice site selection in malignancies.

Herein, we found that the extensive upregulation of spliceosome components is a molecular feature of HCC. Using conditional knock-in mouse as an *in vivo* model, we demonstrated that USP39 was a driver of liver cancer as well as a potent regulator of AS. The splicing landscape regulated by USP39 has been comprehensively characterized in human HCC cells, and KANK2 is one of the functionally important AS targets. Mechanistically, USP39 modulates alternative splice site choice by interacting with SRSF6 and HNRNPC in a conserved manner between humans and mice. USP39 serves as a paradigm of the mid- or late-acting spliceosome components that display potent regulatory properties of AS. These spliceosomal proteins represent potential targets of STTs.

## Results

### USP39-overexpression associates with HCC pathogenesis and aberrant cell cycle signaling

Recent high-throughput technologies have revealed the molecular landscape of HCC. Using a proteomic dataset from CPTAC, we performed unsupervised clustering based on proteins differentially expressed between tumor (T) and non-tumor (NT) livers. Consistent with a previous study(Gao et al., 2019), three subgroups were identified among the 159 tumors. Subgroup 1 was enriched with metabolism-related proteins and subgroup 2 was characterized by immune deregulation. Subgroup 3, with the worst prognosis as reported by Gao et al(Gao et al., 2019), harbored the highest cell renewal pathways, such as cell cycle, DNA replication and spliceosome (Fig. 1A), suggesting that spliceosome proteins may be drivers of HCC subgroup.

**Fig. 1.**
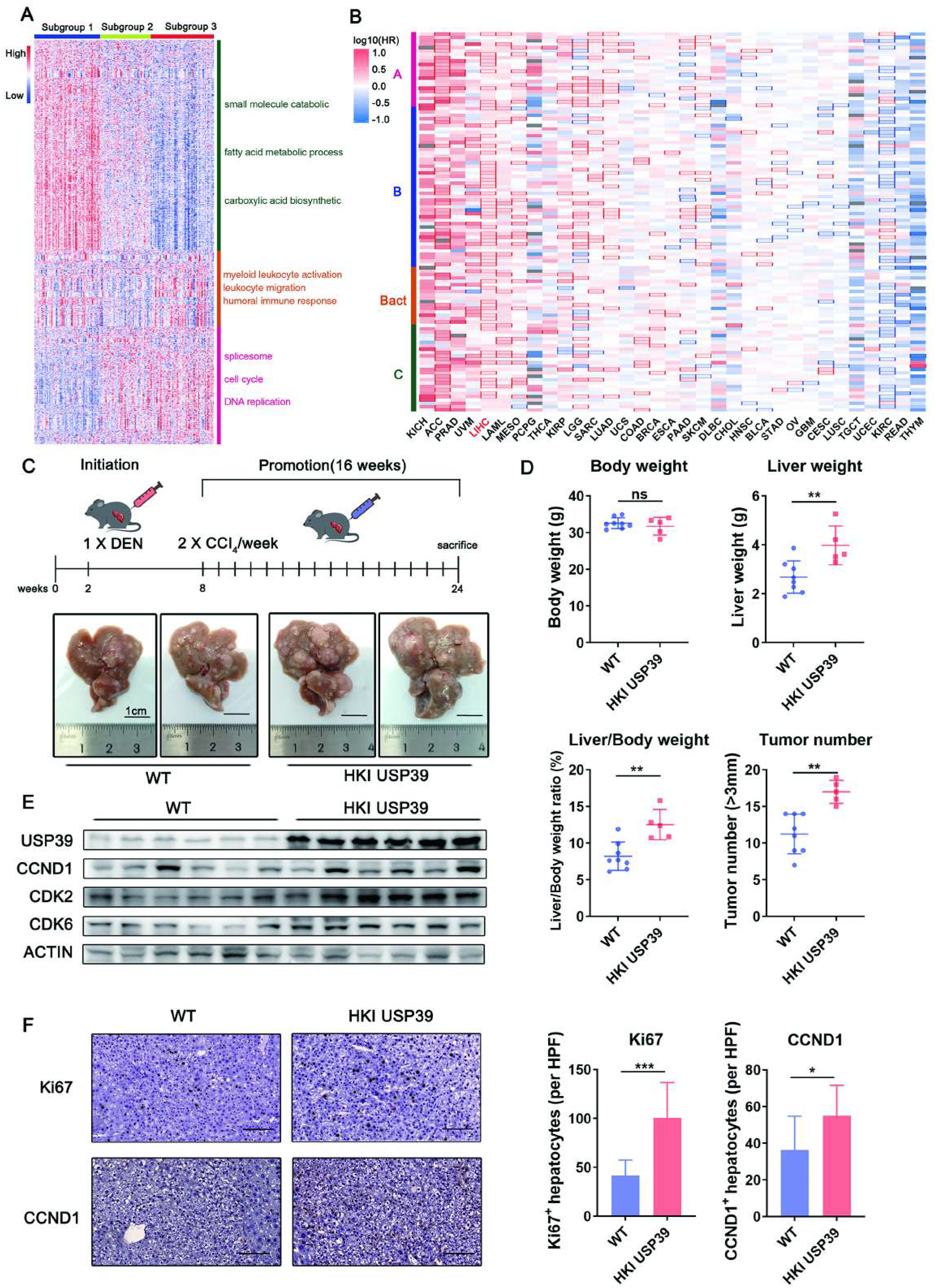
Overexpression of spliceosome components associates with HCC pathogenesis and conditional USP39 knock-in enhances in vivo hepatocarcinogenesis. (A) Proteomic subgroups were identified based on differentially expressed proteins between tumor and non-tumor tissues from the CPTAC HCC cohort. Each column represents a patient sample and rows indicate proteins. The color of each cell shows the Z-score (log2 of relative abundance scaled by protein SD) of the protein in that sample. (B) HR heatmap representing associations between spliceosomal gene expression and OS across 33 different cancer types profiled using TCGA. The HR for the high expression group versus the low expression group (median expression value as a cut-off) was calculated. The heatmap is colored based on the log10 HR. A square with bold border represents a p value < 0.05 in the survival analysis. Each column represents a cancer type, and rows indicate genes arranged in the order of spliceosome assembly (A: complex A, B: complex B, Bact: complex B catalytic active, C: complex C). (C) Schematic summary of the DEN/CCl4-induced hepatocarcinogenesis model established in hepatocyte-specific USP39HKI and control mice (WT). Representative images of tumor-bearing livers are shown. (D) Body weight, liver weight, liver/body weight ratios and tumor numbers of WT and USP39HKI mice were measured. ns: not significant. (E) USP39 and cell cycle related proteins including CCND1, CDK2 and CDK6 were detected with WB. (F) Immunohistochemistry was performed to examine the expression of Ki67 and CCND1 in WT and USP39HKI mice. The IHC-positive hepatocytes were then counted. Three high-power fields (HPF, ×800 magnification) were obtained from each mouse (n=3). Scale bar, 100 μm. Mean ± SD. P values were determined using unpaired Student’s t-test. ∗P <.05, ∗ ∗P <.01, ∗∗∗P < .001. (Experimental group, n=5; control group, n=8)

The spliceosome is a megadalton complex comprising over 100 proteins.(Seiler, Peng, et al., 2018) We studied the effect of spliceosomal gene expression on overall survival (OS) using TCGA data pan-cancer analysis. The hazard ratio (HR) heatmap results showed that spliceosomal gene expression was a robust biomarker for prognosis in various cancers, including liver hepatocellular carcinoma (LIHC) (Fig. 1B). Among these, core tri-snRNP component *USP39* expression was upregulated in paired and unpaired HCC tissues in both TCGA and GEO cohorts (Fig. 1-figure supplement 1A). High *USP39* expression was associated with adverse clinicopathological features and poor prognosis (Fig. 1-figure supplement 1C-D). Consistent with the proteosome results in Fig. 1A, cell cycle and DNA replication pathways were enriched in tumors with high *USP39* mRNA expression via gene set enrichment analysis (GSEA) (Fig. 1-figure supplement 1E). These results highlight the link between USP39 overexpression and HCC pathogenesis, together with changes in cell cycle signaling.

### Conditional USP39 knock-in enhances *in vivo* hepatocarcinogenesis

The clinical data analysis revealed extensive deregulation of spliceosomal genes during HCC development. Whether USP39 serves as a functionally important driver or just a passenger remains to be determined. To establish the causal relationship between USP39 upregulation and hepatocarcinogenesis, hepatocyte-specific USP39 knock-in (USP39HKI) mice were generated by crossing *Rosa26- stopflox/flox-Usp39* mice with *Albumin-Cre* mice. In the DEN/CCl4-induced hepatocarcinogenesis model, we observed more severe liver tumorigenesis in USP39HKI mice than wild-type (WT) mice (Fig. 1C-D). Along with these macroscopic observations, the molecular levels of CCND1, CDK2, CDK6 and Ki67, proteins that promote and mark cell cycle progression, were also upregulated (Fig. 1E-F).

### USP39 extensively modulates alternative splicing through potential interactions with RBPs

To further examine how USP39 promotes tumorigenesis *in vivo*, tumor and para-tumor tissues collected from USP39HKI and WT mice were analyzed by transcriptome sequencing. Successful overexpression of Usp39 was verified, and principal component analysis (PCA) separated tumors from non-tumor tissues (Fig. 2-figure supplement 2). Tumorigenic-related differentially expressed genes (DEGs) and differential alternative splicing (DAS) events were identified by comparing wild-type tumors with wild-type non-tumors (WT-T vs. WT-NT) (Fig. 2A). USP39-regulated events were determined by comparing knock-in non-tumors with wild-type non-tumors (KI-NT vs. WT-NT) (Fig. 2A). DEGs and DAS events regulated by USP39 (KI-NT vs. WT-NT) largely overlapped and positively correlated with tumorigenic events (WT-T vs. WT-NT), indicating USP39 overexpression promotes neoplastic transformation of hepatocytes (Fig. 2B, D). These intersecting DEGs and DAS genes were enriched in cell cycle control, cell proliferation, DNA repair and nuclear division (Fig. 2C, E). Surprisingly, USP39 extensively modulate tumorigenic AS events (2833 out of 8411 events), to a greater extent than its transcriptional regulation of tumorigenic DEGs (382 out of 1799 events). We questioned how this mid/late-acting spliceosome component influence splice site choices. Unlike splicing factors, such as SR and hnRNP proteins, USP39 itself does not possess an RNA-binding domain. We speculate that the specificity of USP39-regualted AS relies on the interplay of USP39 with RBPs, which facilitate the interaction of the spliceosome with corresponding RBP-binding motifs in pre-mRNAs. RBP motif enrichment analysis was performed on overlapping skip exon (SE) events of the KI-NT vs. WT-NT and WT-T vs. WT-NT groups (921 events). A predominant enrichment or depletion of the putative binding motifs was identified within the cassette exons over background exons, implying that USP39 may form regulatory network with RBPs (Fig. 2F, G). In contrast to standard models of AS regulation at early stage of spliceosome assembly, these results provide *in vivo* evidence that the mid/late-acting spliceosomal component USP39 can potently affect splice site selection, possibly via interaction with RBPs.

**Fig. 2.**
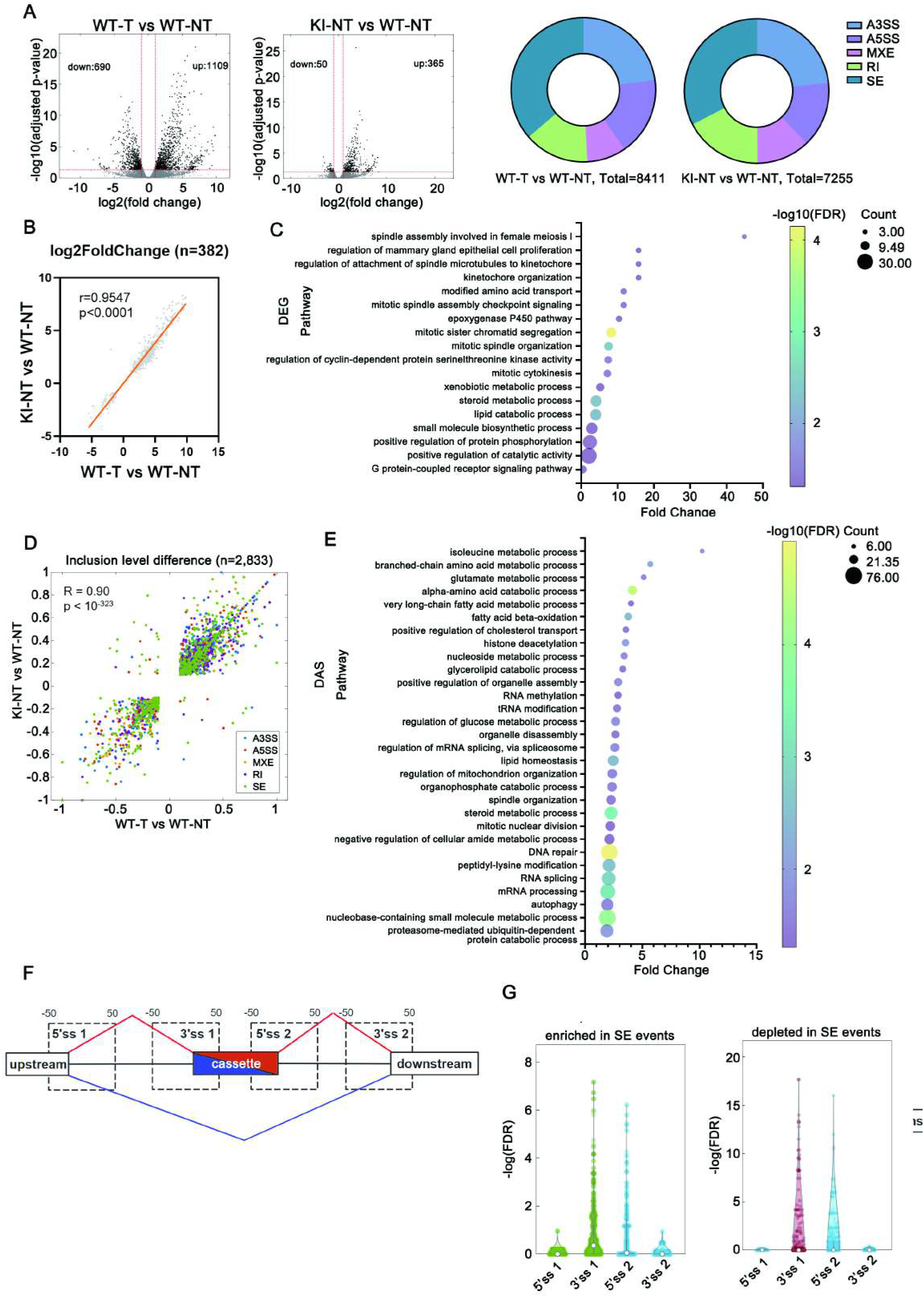
USP39 extensively modulates alternative splicing in mice through potential interactions with RBPs. (A) Volcano plot of DEGs in the WT-T vs. WT-NT and KI-NT vs. WT-NT groups (left). Pie chart depicting the proportions of DAS event types in the WT-T vs. WT- NT and KI-NT vs. WT-NT groups (right). (B) The overlapping DEGs (n=382) were positively correlated in KI-NT vs WT-NT and WT-T vs WT-NT datasets. (C) The GO biological process (BP) enrichment analysis of the overlapping DEGs in KI-NT vs WT-NT and WT-T vs WT-NT datasets. (D) The overlapping DAS events were positively correlated in the KI-NT vs. WT- NT and WT-T vs. WT-NT datasets. (E) The GO biological process (BP) enrichment analysis of the overlapping DAS events in KI-NT vs WT-NT and WT-T vs WT-NT datasets. (F, G) RBP motif enrichment analyses within the 4 regions (5’ss1, 3’ss1, 5’ss2, 3’ss2) around the regulated cassette exons is compared with that around the background cassette exons. The start and end of each region is labelled by a nucleotide distance to the splice site. Red represents shUSP39-activated exons and blue represents shUSP39-repressedd exons. (F) Histogram describing the enriched (left) or depleted (right) motifs in the indicated regions. (G)

### USP39 exerts its pro-proliferative effect in a spliceosome-dependent manner

Consistent with its phenotype in mouse model, USP39 displayed oncogenic functions in human HCC cell lines *PLC-8024* cells and *SNU-449*. ShRNA-mediated USP39 silencing reduced cell viability, foci formation frequencies, colony formation in soft agar and cell cycle progression, whereas USP39 overexpression increased these capabilities (Fig. 3-figure supplement 1-2). It has been reported that USP39 acts as a deubiquitinase as well as a spliceosome component. We wondered whether USP39 promotes HCC cell proliferation through regulating spliceosome assembly. A previous study showed that a C63A mutation in Sad1, the homolog of USP39 in yeast, disrupts its interaction with Snu114 and Snu66, thereby impairing tri-snRNP formation.(Y. H. Huang, Chung, Kao, Kao, & Cheng, 2014) As the core spliceosome component USP39 is highly conserved across species, a USP39-C139A mutant, which is equivalent to the Sad1-C63A mutant, was introduced into USP39-knockdown HCC cells (Fig. 3A, Fig. 3-figure supplement 3A). This point mutation was validated to interrupt tri-snRNP assembly in human HCC cells. RNA immunoprecipitation (RIP) results showed that the decreased U4 and U6 snRNAs were co-purified with the C139A mutant, which indicated specific defects in tri-snRNP biogenesis (Fig. 3B, Fig. 3-figure supplement 3B). Unlike wild-type USP39, which rescued the inhibitory effect of USP39 deficiency on cell proliferation and cell cycle progression, C139A mutant could not reverse the suppressive effects (Fig. 3C-F, Fig. 3-figure supplement 3C-E). These results suggest that USP39 exerts its pro-proliferative effects in a spliceosome-dependent manner.

**Fig. 3.**
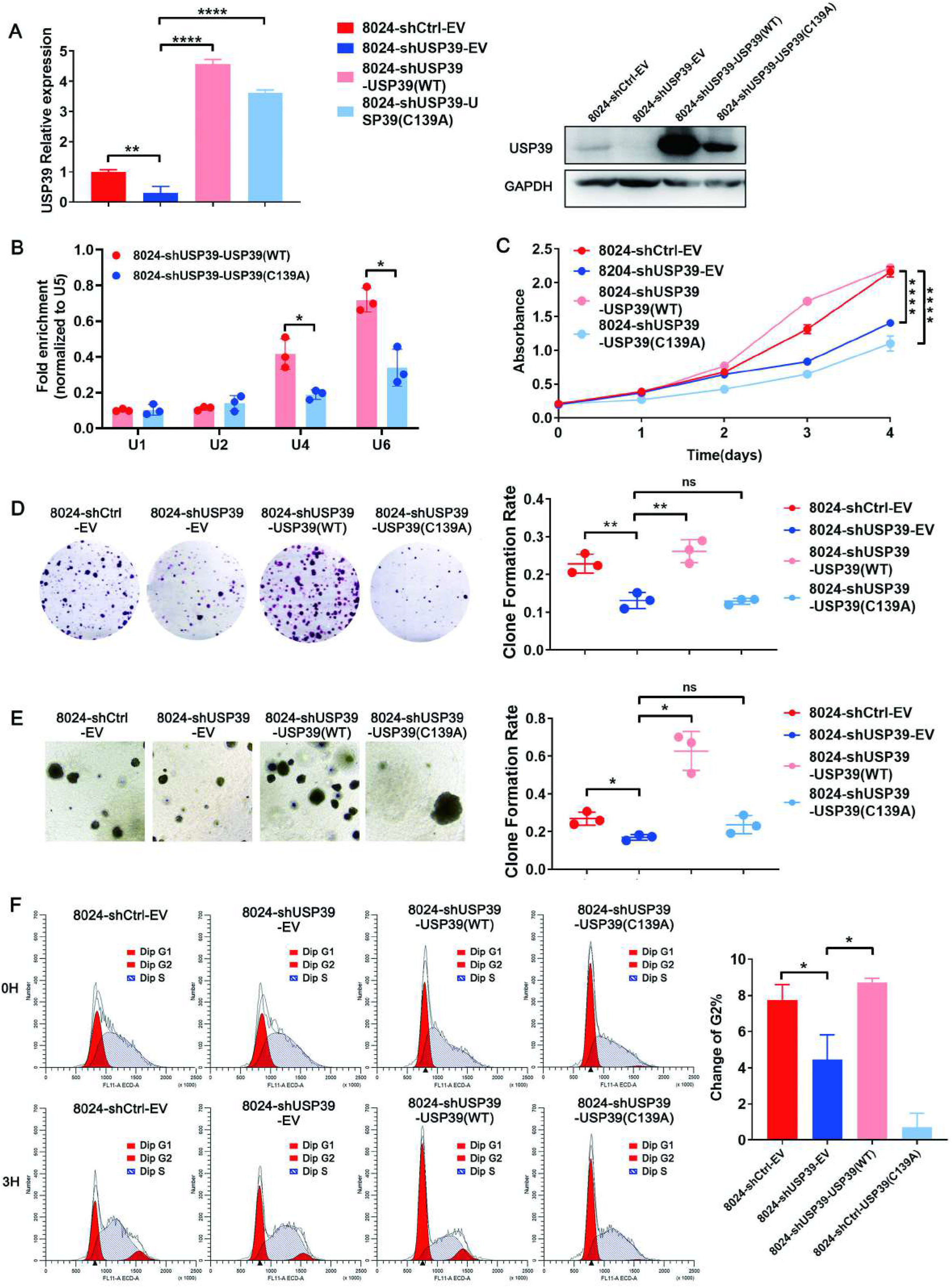
USP39 exerts its pro-proliferative effect in a spliceosome-dependent manner. (A) Flag-USP39 (WT) and Flag-USP39 (C139A) were introduced to USP39- deficiency (shUSP39) PLC-8024 cells and verified with qRT-PCR and WB assays. (B) RIP experiments were performed using antibodies against Flag-USP39 in the indicated cells. The levels of U1, U2, U4, U6 snRNA in immunoprecipitated RNA were detected using qRT-PCR and normalized to the levels of U5 snRNA in each sample (n=3). (C) CCK8 assay showed that only wild-type USP39, but not C139A mutant, could rescue the inhibition effect of USP39 deficiency on cell proliferation. (D, E) Representative images and quantification of foci formation (D) or clone formation in soft agar (E) induced by the indicated cells (n=3). (F) Cells were synchronized at G1/S boundary and detected with PI-stained flow cytometry after release. Cell cycle profiles showed G2 cell populations at 3 hours post-release (n=3). Mean ± SD. P values by paired Student’s t-test (B), or unpaired Student’s t-test (A, C-F). ∗P <.05, ∗∗P <.01, ∗∗∗∗P < .0001. ns: not significant.

### USP39-knockdown reduces global splicing efficiency and selectively regulates AS in human HCC

To verify whether USP39 also regulates human AS, *PLC-8024* USP39-knockdown and control cells were collected for transcriptome sequencing. As shown in Fig. 4A, two independent shRNAs (sh1 and sh2) targeting USP39 generated highly correlated DEG patterns, confirming the reliability of the RNA-seq data. Gene ontology (GO) enrichment analysis of these overlapping DEGs showed that USP39 depletion significantly affected pathways such as cell adhesion, translation, and cell cycle (Fig. 4B). As USP39 is required for spliceosome complex B assembly and catalytic activation, we investigated whether *USP39* expression affects pre-mRNA splicing. We calculated splicing efficiency using RNA-seq data and found a widespread impairment in pre-mRNA splicing upon the loss of USP39. Both 5’ and 3’splice site efficiencies were significantly reduced (Fig. 4C), suggesting that USP39 acts as a global regulator of splicing efficiency. DAS events (Δ percent spliced in, ΔPSI> 0.1, and adjusted p-value<0.05) were further analyzed, and 522 overlapping events were detected in sh1 and sh2 knockdown cells. These two USP39 deficient cells shared strongly correlated DAS patterns (correlation coefficient r = 0.93), which ensured data reliability (Fig. 4D). Among these 522 events, 349 were skipped exon (SE) events with either increased or decreased PSI values (Fig. 4E). Five target genes that have been reported to influence cell viability or cell cycle transition were selected and verified by RT-PCR (Fig. 4F).

**Fig. 4.**
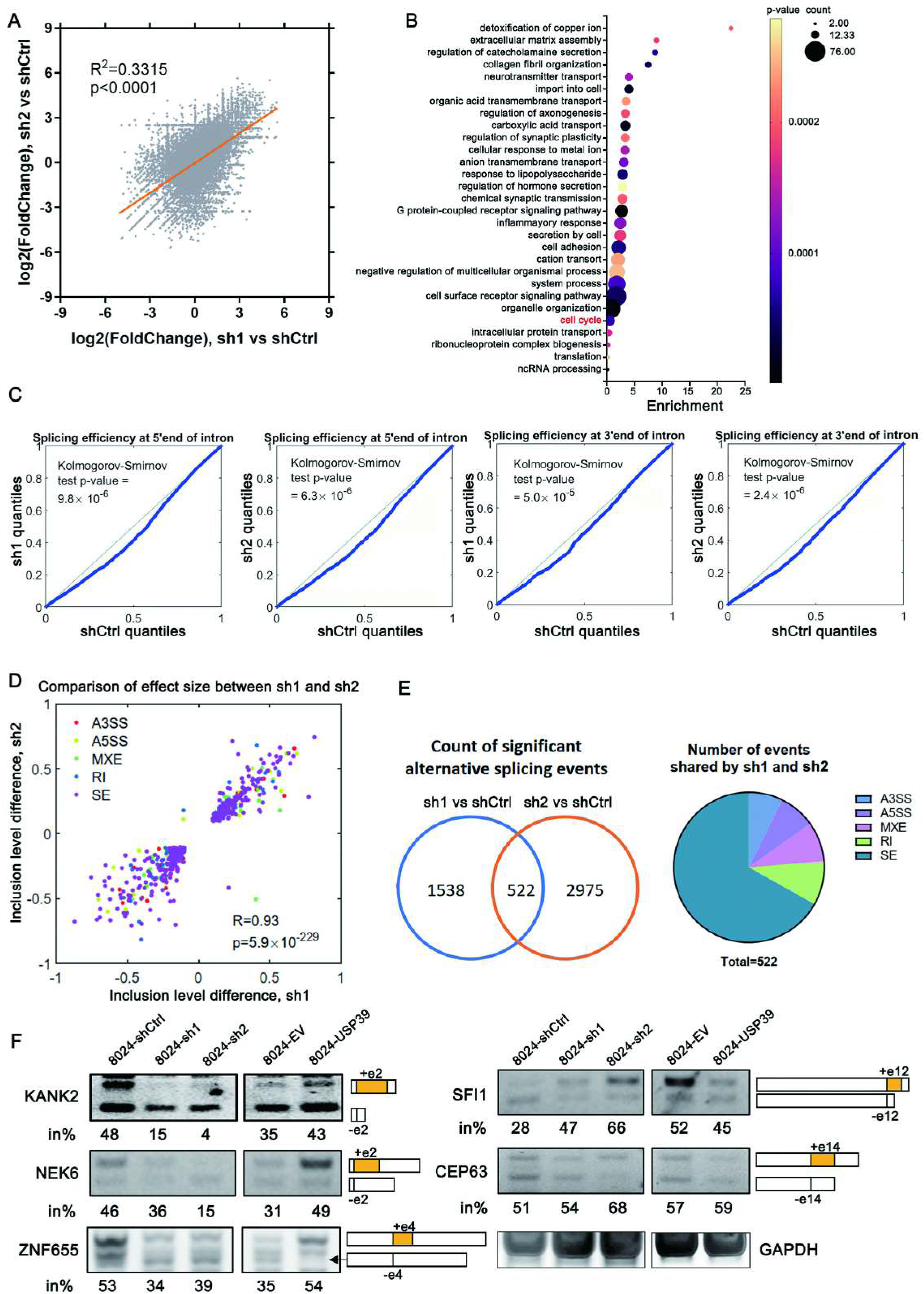
USP39 knockdown reduces global splicing efficiency and selectively regulates alternative splicing. (A) PLC-8024 USP39-knockdown (sh1, sh2) and control (shCtrl) cells were collected for transcriptome sequencing. Sh1 showed similar DEG pattern to sh2. (B) The GO biological process (BP) enrichment analysis of the overlapping DEGs. (C) Global splicing efficiency analysis at 5′ and 3′ splicing sites in PLC-8024 cells with and without USP39 depletion. (D) DAS events were highly correlated in sh1 vs. shCtrl and sh2 vs. shCtrl datasets. (E) Venn diagram displayed the DAS events in two datasets after the intersection (left). Pie chart depicting the proportions of different types of overlapping DAS events (right). A5SS: alternative 5’ splice site, A3SS: alternative 3’ splice site, MXE: mutually exclusive exons, RI: retained intron, SE: skipped exon. (F) Three events with down-regulated PSI values (left) and two targets with up- regulated PSI values (right) in USP39 depletion PLC-8024 cells were validated with RT-PCR and agarose gel electrophoresis. The percentages of cassette exon inclusion over the total transcripts are presented using in%.

### USP39 promotes tumorigenesis partially through a splicing switch of KANK2

Among the USP39-affected DAS events, we focused on the KN motif and ankyrin repeat domains 2 (KANK2) gene, a focal adhesion protein that regulates cell adhesion, migration, apoptosis, and proliferation.(Gee et al., 2015; Pei et al., 2022; D. Wang et al., 2012) Knockdown of USP39 resulted in a decrease in the long isoform of KANK2 (KANK2-L, containing exon 2) relative to its short isoform (KANK2-S, lacking exon 2), whereas USP39 overexpression caused opposite changes (Fig. 4F). Based on these *in vitro* findings, we first examined the clinical significance of these two KANK2 isoforms by comparing the proportion of KANK2- L mRNA in 66 human HCC and adjacent non-tumor tissues. Both the expression of USP39 and the inclusion of KANK2 exon 2 were significantly increased in tumor samples compared to non-tumor samples in this in-house HCC cohort (Fig. 5A). A positive correlation was observed between *USP39* mRNA expression and the percentage of KANK2-L (Fig. 5B). Patients with either higher levels of *USP39* expression or a higher proportion of the KANK2-L isoform had poorer prognosis, although this was not statistically significant due to the limited number of cases. Notably, patients with relatively high *USP39* expression and a high proportion of KANK2-L showed significantly worse OS than those with relatively low *USP39* expression and a low proportion of KANK2-L (Fig. 5C).

**Fig. 5.**
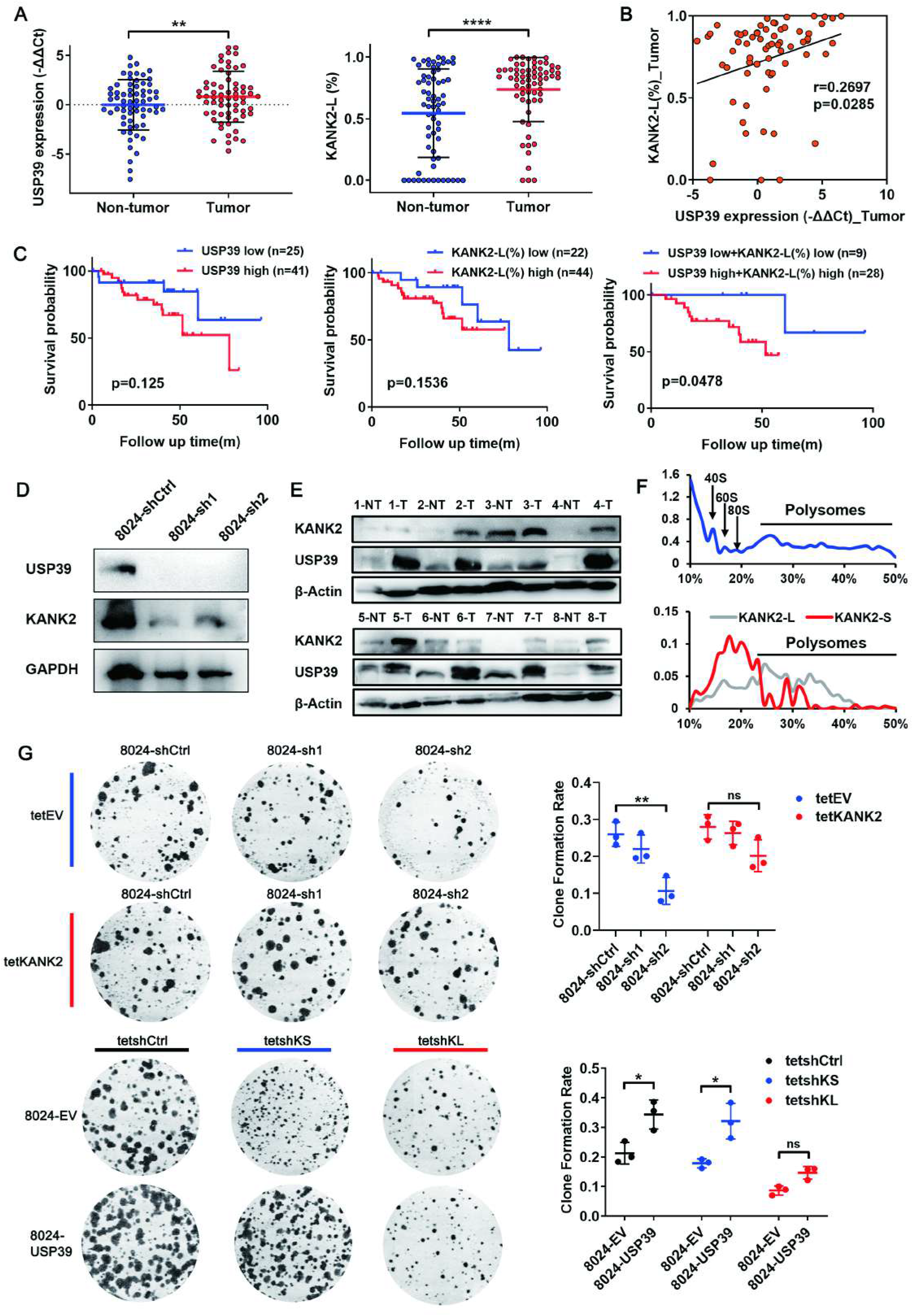
USP39 exerts its malignant effects partially through an isoform switch of KANK2-S to KANK2-L. (A) The mRNA level of USP39 and the expression proportion of KANK2-L isoform were detected in our in-house HCC cohort (n=66). (B) The expression proportion of KANK2-L isoform was positively correlated with USP39 mRNA expression in tumor tissues. (C) Kaplan–Meier OS curves of in-house HCC patients based on USP39 expression (high vs. low, defined as tumor/non-tumor > 1) and KANK2-L percentage (high vs. low, defined as tumor(%)/non-tumor(%)>1). (D) WB assay revealed KANK2 protein reduction in USP39 knockdown cells. (E) WB analysis of adjacent non-tumor (NT) and tumor (T) tissue lysates from HCC patients with USP39 and KANK2 antibodies. β-Actin was used as a loading control. (F) Polyribosome profile analysis revealed that KANK2-L isoform enhanced the translation of KANK2 protein. (G) Tet-KANK2 was introduced to USP39-deficient PLC-8024 cells (up). Two Tet- On shRNAs targeting KANK2-L (tetshKL) and KANK2-S (tetshKS) were introduced into USP39-overexpressing PLC-8024 cells (down). Foci formation assay was performed to access growth of the indicated cells. Representative images and statistical results are shown (n=3). Mean ± SD. p values by unpaired Student’s t-test. ∗P < .05, ∗∗P < .01, ns: not significant.

We examined the molecular functions of the two KANK2 isoforms. Exon 2 is located in the 5’-untranslated region (UTR) of KANK2 and does not affect the protein sequence. WB of KANK2 protein showed only one band, but a marked reduction in KANK2 protein was observed in accordance with the isoform switch in USP39 knockdown cells (Fig. 5D). Furthermore, a positive correlation was observed between USP39 and KANK2 protein expression in HCC clinical samples (Fig. 5E). Therefore, we hypothesized that the KANK2-L isoform enhances the translation of the KANK2 protein compared to KANK2-S. Supporting this, polyribosome profile analysis revealed that a higher proportion of KANK2-L was associated with heavy polyribosomes, whereas KANK2-S was largely associated with light polyribosomes (Fig. 5F). Mass spectrometry analysis of RNA pulldown samples showed that KANK2-L mRNA bound more translation related proteins than KANK2-S mRNA, indicating a higher translation efficiency for KANK2-L (Fig. 5-figure supplement 1A). By applying the Tet-On system to replenish KANK2 expression upon doxycycline induction, we found that restoration of KANK2 partially reversed the adverse effects of USP39 knockdown on HCC cell proliferation and cell cycle control (Fig. 5G, Fig. 5-figure supplement 1B, D, and F), underscoring the importance of USP39-regulated KANK2 splicing in hepatocarcinogenesis. To provide further evidence, two Tet-On shRNAs specifically targeting KANK2-L and KANK2-S were introduced into USP39- overexpressing HCC cells. KANK2-L depletion markedly suppressed the pro- proliferation phenotype of USP39, whereas KANK2-S silencing had minor effects (Fig. 5G, Fig. 5-figure supplement 1C, E and F). WB analysis confirmed that KANK2-L depletion correspondingly reduced KANK2 protein expression, while KANK2-S targeting barely affected KANK2 protein expression, despite successful knockdown of KANK2-S mRNA (Fig. 5-figure supplement 2). These data suggest that USP39 exerts its malignant effects partially through an isoform switch from KANK2-S to KANK2-L, with a shift in translation of KANK2 from low efficiency to high efficiency.

### USP39 selectively regulates exon inclusion/exclusion via interaction with SRSF6/HNRNPC in a position-dependent manner

Data from the mouse model suggested that USP39 modulated AS by forming a regulatory network with RBPs. We continued to perform RBP motif enrichment analysis on shUSP39-regulated exon skip events in human *PLC-8024* HCC cells. RBP-binding motifs maintained a strictly similar distribution pattern in humans and mice, including the depleted SRSF6-binding motif and the enriched HNRNPC-binding motif (Fig. 6A, Supplementary file 1).

**Fig. 6.**
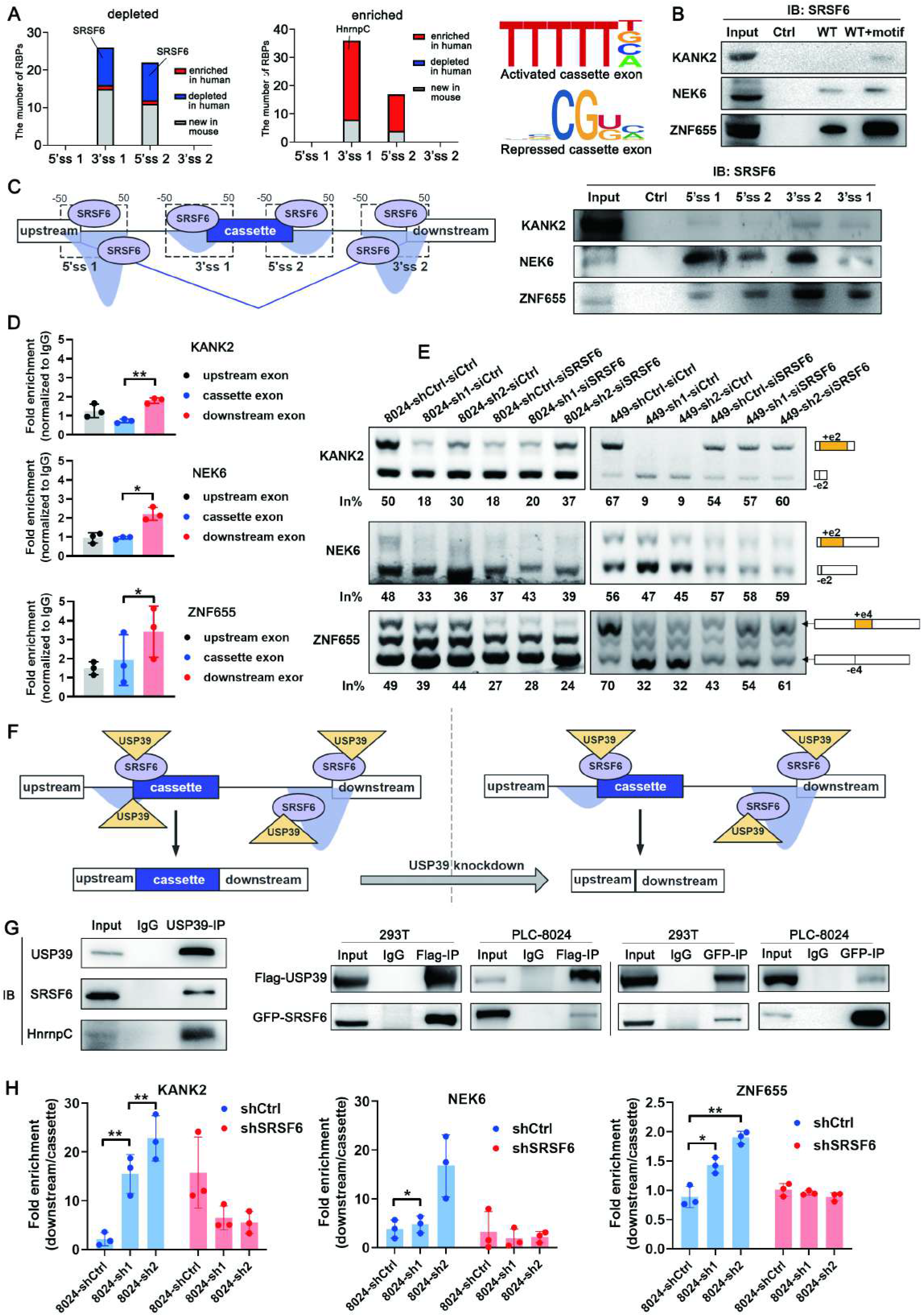
USP39 selectively regulates exon inclusion/exclusion via interaction with SRSF6 in a position-dependent manner. (A) Comparison of motif enrichment analysis results between human HCC cells and mice. The depleted SRSF6-binding motif and the enriched HNRNPC-binding motif were distributed in a conserve pattern in humans and mice. (B, C) RNA pulldown assay verified the binding of SRSF6 to the putative binding motif (B) and predominant binding of SRSF6 within the flanking constitutive exons over the cassette exons (C). Representative images of three independent reproducible experiments are shown. (D) RIP assay was performed in PLC-8024 cells with SRSF6 antibody or IgG control (n=3). (E) Minigene reporters of KANK2, ZNF655 and NEK6 were introduced into USP39- knockdown PLC-8024 and SNU-449 cells. SRSF6 was further silenced and splicing of the minigenes was verified by RT-PCR. The percentages of cassette exon inclusion within the total transcripts are presented using in%. (F) Schematic illustration of the mechanism by which shUSP39 represses exon inclusion through interacting with SRSF6 in a position-dependent manner. (G) Endogenous (left)/ ectopic overexpressed (right) USP39 was coimmunoprecipitated with endogenous (left)/ ectopic overexpressed (right) SRSF6 protein. (H) RIP assay showed that USP39 retained in the flanking constitutive exons upon USP39 knockdown (blue columns). Silencing of SRSF6 in USP39-knockdown cells abolished USP39 retainment in the flanking constitutive exons (red columns). Data were presented as fold enrichment between downstream exon signal and cassette exon signal (downstream/cassette). Mean ± SD. P values were determined using unpaired Student’s t-test. ∗P <.05, ∗ ∗P <.01.

The putative SRSF6-binding motif was among the top predicted motifs derived from the shUSP39-repressed exon set. The above-mentioned three shUSP39- repressed AS events (KANK2, NEK6, and ZNF655) all contained SRSF6-binding motifs in the flanking constitutive exon junction, but not in the cassette exon junction. Therefore, we investigated whether SRSF6 participates in shUSP39- repressed exon splicing. To this end, an RNA pulldown assay was performed to examine whether SRSF6 binds to this putative binding site. Biotin-labeled cassette exon RNAs (WT) were transcribed *in vitro* and the putative SRSF6-binding motif was inserted into these RNAs (WT+motif). Compared with WT RNAs, WT+motif RNAs pulled down more SRSF6 protein in all three tested genes (Fig. 6B), indicating that SRSF6 can recognize and bind to this putative SRSF6-binding motif. Thereafter, these cassette exons together with the upstream and downstream constitutive exons were compared for their SRSF6 binding affinities using an RNA pulldown assay. The SRSF6 binding pattern was consistent with the motif distribution feature of shUSP39-repressed exons, which was characterized by the predominant binding of SRSF6 within the flanking constitutive exons over the cassette exons (Fig. 6C). Consistently, the RIP assay showed enrichment of SRSF6-bound flanking constitutive exon RNAs over cassette exon RNAs (Fig. 6D). To gain mechanistic insights into shUSP39-repressed exon inclusion, we constructed minigene reporters of KANK2 exons 1-3, NEK6 exons 1-3 and ZNF655 exons 3-5 respectively. Splicing was assayed following transient transfection of the control and USP39-knockdown sub–cell lines. In accordance with the endogenous splicing pattern (Fig. 4F), the inclusion of KANK2 exon 2, NEK6 exon 2, and ZNF655 exon 4 was markedly inhibited by USP39 knockdown (Fig. 6E, lanes 1-3, 7-9). On this basis, siRNA-mediated SRSF6 silencing substantially restored the inclusion of cassette exons in the USP39-knocdown sub- cell line (lanes 5-6, 11-12) to a level equivalent to that in the control sub-cell line (lanes 4 and 10), indicating that the shUSP39-repressed inclusion of KANK2, NEK6, and ZNF655 cassette exons was SRSF6 dependent. Based on the above results, we proposed a model in which the enrichment of SRSF6-binding motifs within the flanking constitutive exons recruits SRSF6 and subsequently detains relatively more USP39 during USP39 knockdown, resulting in repressed cassette exon inclusion (Fig. 6F). Supporting this, SRSF6 protein was coimmunoprecipitated with USP39 in human embryonic kidney *293T* cells and *PLC-8024* HCC cells, suggesting that SRSF6 can interact and help recruit USP39 (Fig. 6G). This binding between SRSF6 and USP39 aids the recruitment of USP39 to the SRSF6-binding motif-enriched region. As a result, RIP analysis showed that USP39 was retained in the flanking constitutive exons upon USP39 knockdown, whereas SRSF6 silencing abolished USP39 retention (Fig. 6H).

Similarly, we investigated the mechanism by which shUSP39 activates cassette exon inclusion. The motifs derived from the shUSP39-activated exons indicated a predominant enrichment of HNRNPC binding motifs within the cassette exons over the flanking constitutive exons. Putative HNRNPC-binding motif and two shUSP39-activated AS events (SFI1 and CEP63) were selected to demonstrate the underlying mechanism. The RNA pulldown assay confirmed binding between HNRNPC and the putative binding motif (WT), which was almost abolished when the motif was deleted (Δmotif Mut)(Fig. 7A). Both the RNA-bound HNRNPC and HNRNPC-bound RNA patterns were consistent with the motif distribution features of the shUSP39-activated exons (Fig. 7B, C). Minigene reporter assays showed that the shUSP39-activated inclusion of SFI1 and CEP63 cassette exons was HNRNPC-dependent (Fig. 7D). Based on the above results, we propose a model in which the enrichment of HNRNPC-binding motifs within the cassette exons recruits HNRNPC and subsequently detains relatively more USP39 during USP39 knockdown, which results in activated cassette exon inclusion (Fig. 7E). Supporting this, endogenous/ectopically overexpressed USP39 was coimmunoprecipitated with endogenous/ectopic overexpressed HNRNPC protein, suggesting that HNRNPC can interact and help recruit USP39 (Fig. 6G, 7F). RIP analysis confirmed that USP39 was retained in HNRNPC-binding motif-enriched cassette exons upon USP39 knockdown, which was abrogated after HNRNPC silencing (Fig. 7G). Moreover, the Co-IP assay also detected interaction between USP39 and SRSF6/HNRNPC in various mouse cell lines, suggesting this regulatory network in conserved between humans and mice (Fig. 7H). Collectively, these results demonstrated that USP39 selectively regulates exon inclusion/exclusion via interaction with SRSF6 or HNRNPC in a position- dependent manner.

**Fig. 7.**
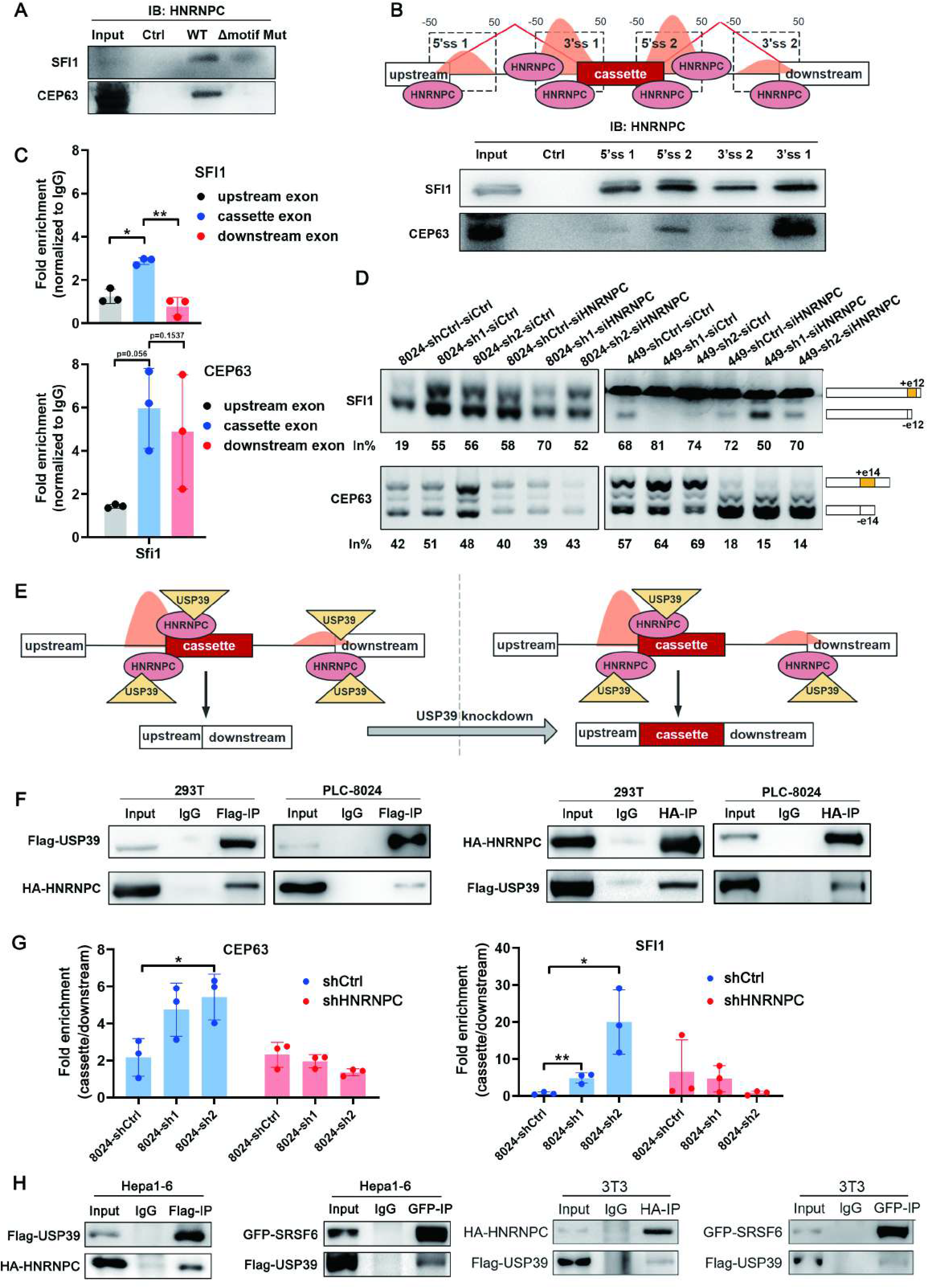
USP39 selectively regulates exon inclusion/exclusion via interaction with HNRNPC in a position-dependent manner. (A, B) RNA pulldown assay verified the binding of HNRNPC to the putative binding motif (A) and predominant binding of HNRNPC within the cassette exons over the flanking constitutive exons (B). Representative images of three independent reproducible experiments are shown. (C) RIP assay was performed in PLC-8024 cells using antibody against HNRNPC or IgG control. The levels of precipitated cassette exon RNA and the flanking exon RNA were detected using qRT-PCR and the data were presented as fold enrichment relative to IgG control (n=3). (D) Minigene reporters of SFI1 and CEP63 were introduced intro USP39- knockdown PLC-8024 and SNU-449 cells. HNRNPC was further silenced and splicing of the reporters was verified by RT-PCR. The percentages of cassette exon inclusion within the total transcripts are presented using in%. (E) Schematic illustration of the mechanism by which shUSP39 activates exon inclusion through interacting with HNRNPC in a position-dependent manner. (F) 293T and PLC-8024 cells were co-transfected with a combination of two plasmids expressing Flag-USP39 and HA-HNRNPC proteins. co-IP experiments with anti-Flag or anti-HA agarose antibodies and visualized by WB analysis using anti-Flag and anti-HA antibodies, respectively. (G) RIP assay using USP39 antibody showed that USP39 retained in the cassette exons upon USP39 knockdown (blue columns). Silencing of HNRNPC in USP39- knockdown cells abolished USP39 retainment in the cassette exons (red columns). Data were presented as fold enrichment between cassette exon signal and downstream exon signal (cassette/downstream). (H) Ectopic overexpressed Flag-USP39 was coimmunoprecipitated with ectopic overexpressed HA-HNRNPC/GFP-SRSF6 proteins in Hepa1-6 and 3T3 cells. Mean ± SD. P values by unpaired Student’s t test. ∗P <.05, ∗∗P <.01.

## Discussion

Spliceosome is one of the most complex molecular machineries of the cell. At present, models of splicing are mainly delineated by detailed biochemical studies using yeast or a small number of model introns.(Papasaikas, Tejedor, Vigevani, & Valcárcel, 2015) Considering the close relationship between AS and diseases including cancer, *in vivo* models and data are especially needed to refine the existing regulatory networks. In the current study, we profiled the global AS landscape in wildtype and USP39 conditional knock-in mouse HCC. Knock-in of USP39 simulates its overexpression in human HCC and causes AS changes of higher magnitude than gene expression changes in mice. This great effect on splice site selection implies that the standard models of AS regulation may underweight contributions from mid- or late-acting spliceosome proteins. Data from human HCC cells *in vitro* also support this notion by providing more detailed biochemical evidence. Although USP39 has been demonstrated to regulate cell activities via different mechanisms, including changing constitutive splicing efficiency and deubiquitinating protein substrates.(L. Dong et al., 2020; Liao, Li, Liu, & Song, 2021; Ni et al., 2021; Pan et al., 2015; Yuan et al., 2015) We found that USP39 promoted hepatocarcinogenesis in a spliceosome-dependent manner. Supporting this, only WT USP39, but not C139A USP39 (a mutant that impairs spliceosome assembly), rescued the malignant phenotypes of USP39-silenced HCC cell lines. Moreover, DAS events affected by USP39 participate in a wide range of tumor-related functions such as cell cycle control. Splicing switch of *KANK2* has been demonstrated to be an important molecule that mediates the oncogenic effects of USP39 in HCC. These findings suggest that mid/late-acting spliceosome component USP39 is a potent regulator of AS and the resulting isoform switching is functionally important. Of note, our data do not rule out other possible mechanisms. In fact, USP39 knockdown also reduces global constitutive splicing efficiency, which is tightly coupled with transcription and gene expression. Previous studies have mainly stated the role of U1 and U2 snRNP components in splicing site selection. However, how can late-acting spliceosome components such as USP39 influence splice site choices? Substantial studies have provided a consensus picture in which splicing factors recognizing cognate auxiliary sequences in the pre-mRNA promote or inhibit early events in spliceosome assembly (e.g., U1 snRNP and U2 snRNP recruitment).(Fu & Ares, 2014) These regulatory factors include members of hnRNP and SR protein families, which often display cooperative or antagonistic functions depending on the position of their binding sites relative to the regulated splice sites.(Barash et al., 2010; Zhou et al., 2019) In our study, we proved that these hnRNP and SR proteins also regulated the recruitment of U4/U6. U5 tri-snRNP components, thereby influencing specific splicing site recognition. Based on the mechanistic investigations, including motif analyses, RNA pulldown assays, RIP, and minigene reporter assays, we found that depletion of SRSF6-binding motifs in the cassette exons always led to exon exclusion, whereas enrichment of HNRNPC-binding motifs within the cassette exons always led to exon inclusion upon USP39 knockdown. Intriguingly, this mechanism is conserved in mice and humans. Although a considerable fraction of AS events appears to be species-specific, the interaction between USP39 and SRSF6/HNRNPC is conserved between humans and mice. It is conceivable that such RBPs would favor the splicing of alternative splice sites they recognize by recruiting limited amounts of the spliceosome component USP39. In this context, regulatory plasticity may be contributed, at least in part, to the substantial number of splicing factors containing disordered regions. USP39, SRSF6, and HNRNPC contain intrinsically disordered regions.(Geuens, Bouhy, & Timmerman, 2016; Hadjivassiliou, Rosenberg, & Guthrie, 2014; Zheng et al., 2020) Such regions may be flexible to adopt different conformations, allowing alternative routes for spliceosome assembly on different introns, with some splicing sites being more sensitive than others to the depletion of a core spliceosomal factor.(Shepard & Hertel, 2009)

Besides SRSF6 and HNRNPC, RBP-binding motif enrichment analyses also revealed that other RBPs are potentially involved in USP39-modulated splicing. And there are nearly 100 mid- or late-acting spliceosome components in addition to USP39, which reminds us about the extraordinary complexity of AS regulatory networks. The RBP-binding motif enrichment analyses we developed in this study illustrates the potential to identify molecular mechanisms of regulation on the basis of RNA-seq data. There is now a wealth of publicly available RNA-seq datasets generated upon knockdown of hundreds of RNA-related genes as part of the ENCODE project. This may provide abundant bioinformation for us to systematically construct the AS networks.

In summary, the AS landscape profiled in mouse HCC highlights the remarkable regulatory potential of USP39 in splice site selection. USP39 is a driving factor for hepatocarcinogenesis, whose function is carried out in a spliceosome-dependent manner and at least partially mediated by oncogenic splicing switch of *KANK2* gene. Our analysis provided a new regulatory model that is conserved in humans and mice, whereby USP39 regulates exon inclusion/exclusion by interacting with SRSF6 or HNRNPC in a position-dependent manner. These findings refine the standard model by demonstrating mid/late-acting spliceosome components can regulate splicing site selection through cooperation with other splicing regulators. USP39 and possibly other mid/late-acting spliceosome components may serve as tumor biomarkers and therapeutic targets. As a general approach, our data suggest that systematic identification of AS targets can be combined with bioinformatics and biochemistry to identify the sites and mechanisms of action of alternative splicing factors.

## Materials and Methods

### Clinical samples

66 pairs of HCC and para-tumor tissue samples were collected with informed consent from patients who underwent hepatectomy at Sun Yat-Sen University Cancer Center (Guangzhou, China). The clinical specimens used in this study were approved by the Committee for Ethical Review of Research Involving Human Subjects at the Sun Yat-Sen University Cancer Center (GZR2020-260). Complete clinicopathological and follow-up data were available and the patient demographic information is provided in Supplementary file 2.

### Animal experimentation

All animal experiments were reviewed and approved by the Institutional Animal Care and Use Committee, Southern University of Science and Technology (SUSTC-2019-069). All mice had a C57bl/6 background. USP39HKI mice were obtained by crossing *Rosa26-stopflox/flox-USP39* (Cyagen Biosciences, Guangzhou, China) and *Albumin-Cre* (Shanghai Model Organisms Center, Shanghai, China) mice. The resulting *Rosa26-stopflox/flox-Usp39*; *Alb-Cre* mice served as experimental group and *Alb-Cre/+* mice served as control group. Only the male mice were used in this study.

To establish the HCC model, intraperitoneal injection of 20 mg/kg diethylnitrosamine (DEN) (Sigma, #N0258-1g, Missouri, USA) was performed within two weeks after birth to initialize the HCC process. CCl4 (2ul/g body weight, prediluted 1:10 in olive oil) was intraperitoneally injected twice a week after an interval of 6 weeks to promote HCC progression. The mice were sacrificed after 16-week promotion and the liver/body weight, tumor number, and expression levels of CCND1 and Ki67 were assessed.

### RBP motif enrichment analysis

For RBP motif enrichment analysis, background sequences were generated by randomly selecting 150,000 exons from the GRCh38 or GRCm39 reference genome and retrieving DNA sequences of their 5’ and 3’ splice sites (i.e., regions with length of 100bp flanking the 5’ and 3’ ends of these exons). The numbers of RBP motifs in splice sites associated with significant alternative splicing events and the random background sequences were counted using HOMER, and then used in enrichment analysis of these motifs based on one-sided Fisher’s exact- test (left-sided test for identification of deprived motifs and right-sided test for enriched motifs) implemented in MATLAB scripts. P-values were adjusted using the Benjamini-Hochberg procedure for multiple hypothesis testing.

### Consensus clustering for proteomic data

HCC proteomic data were retrieved from the Clinical Proteomic Tumor Analysis Consortium (CPTAC). Consensus clustering was implemented for 1,274 differentially expressed proteins (Gao et al., 2019) using the ConsensusClusterPlus R package, and the following parameters were used for clustering: number of repetitions = 1,000 bootstraps; pItem = 0.8 (resampling 80% of any sample); pFeature = 0.8 (resampling 80% of any protein); and k-means clustering with up to 6 clusters, and 3-cluster as the optimized solution for clustering. Gene Ontology (GO) and Kyoto Encyclopedia of Genes and Genomes (KEGG) enrichment analyses were performed using the online tool MSigDB (https://www.gsea-msigdb.org/gsea/msigdb/).

### Survival-related analyses

Using the GEPIA2 webserver (http://gepia2.cancer-pku.cn/), the association between each spliceosomal gene (Seiler, Peng, et al., 2018) expression level and overall survival (OS) for all cancers was determined by univariate Cox regression for the highly expressed group versus the low one, with the median expression as a cut-off value. The log-rank test was used to compare differences in survival distribution.

### TCGA and GEO data analyses

The HCC transcriptome data were obtained from The Cancer Genome Atlas Liver Hepatocellular Carcinoma (TCGA_LIHC) project and the NCBI Gene Expression Omnibus (GEO) database (accession no. GSE124535, GSE14520). Gene Set Enrichment Analysis (GSEA) (http://www.broadinstitute.org/gsea) was performed to identify associated molecular pathways.

### Western Blot

Samples were lysed with Pierce RIPA buffer (Thermo Scientific, Massachusetts, USA) and quantified using Bio-Rad Protein Assay (Bio-Rad, California, USA). After being mixed with loading dye (Bio-Rad), protein was denatured (100℃, 15min). Denatured proteins were separated by SDS-PAGE and transferred onto a PVDF membrane (Millipore, Massachusetts, USA). After blocking, the membranes were incubated with primary antibodies (USP39, Abcam, Cambridge, UK, #ab131244; CCND1, CST, Massachusetts, USA, #2978; CDK2, CST, #2546; CDK6, CST, #3136; β-actin, CST, #3700; GAPDH, CST, #2118; KANK2, Proteintech, Illinois, USA, # 21733-1-AP; SRSF6, ABclonal, Massachusetts, USA, #A14603; HNRNPC, Proteintech, #11760-1-AP; Flag, Proteintech, #20543-1-AP; GFP, RUIXIN, Quanzhou, China, # GXP204228; HA, CST, #3724) and incubated with peroxidase-conjugated secondary antibodies. Immunoreactive bands were visualized using ECL (Bio-Rad) and exposed to autoradiograph film.

### Immunohistochemistry

Liver tissues were fixed in 4% paraformaldehyde, embedded in paraffin, and sectioned at 5 µm thickness for IHC staining. After blocking with 5% bovine serum albumin, the deparaffinized sections were incubated with primary antibodies (Ki67, CST, #12202; CCND1, CST, #2978) overnight at 4 °C. Antibody binding was detected by incubation with biotinylated anti-rabbit IgG antibody and visualized by reaction with DAB Substrate (Boster, California, USA).

### Cell lines

The HCC cell line *PLC-8024* (TCHu119) was obtained from the Institute of Virology of the Chinese Academy of Sciences (Beijing, China). The HCC cell line *SNU-449* (CRL-2234) was obtained from the American Type Culture Collection (ATCC). Cell line *293T* (SCSP-502) was obtained from the Cell Bank affiliated to Shanghai Institute of Biochemistry and Cell Biology. The mouse HCC cell line *Hepa1-6* (CTCC-ZHYC-0566) was obtained from Meisen Chinese Tissue Culture Collections (MeisenCTCC). The mouse embryo fibroblast cell line *3T3* was a kind gift from Dr. Xijun Ou. All cell lines were authenticated using short tandem repeat profiling and routinely tested for mycoplasma contamination. The cells were cultured in Dulbecco’s modified Eagle’s medium (DMEM) (Gibco, California, USA), supplemented with 10% fetal bovine serum (Gibco), and 1% penicillin/streptomycin mixture (Gibco). All cell lines used in this study were incubated at 37°C in a humidified incubator containing 5% CO2.

### siRNA transfection

siRNA sequences were designed using the siDRECT website (http://sidirect2.rnai.jp/) and the sequences are listed in Supplementary file 3. siRNA transfection was performed using siRNA-mate reagent (GenePharma, Shanghai, China) according to the manufactures’ instruction.

### Plasmids, lentivirus production and cell infection

Wild-type and mutant (C139A) USP39 with a Flag-tag at its N-terminal were cloned into the pLenti6/V5 lentiviral vector (Invitrogen, California, USA). Specific shRNA oligonucleotides targeting USP39 were cloned into the pLL3.7 lentiviral vector (Addgene). These plasmids, together with lentivirus packaging vectors from the pLenti6/V5 Directional TOPO Expression Kit (Invitrogen), were co-transfected into *HEK293T* cells using Lipo3000 (Invitrogen). After 3-day incubation, the medium containing the specific virus was collected and added to *PLC-8024* and *SNU-449* cells combined with 10μg/ml polybrene. After 72 h, pLenti6/V5-Flag-USP39 infected cells were selected with 5 μg/ml blasticidin (Gibco) while pLL3.7-shRNA infected cells were selected with 2μg/ml puromycin (Gibco). The overexpression and knockdown efficiencies were validated with WB and qRT-PCR. primers and USP39-shRNAs sequences are listed in Supplementary file 3.

### Tet-On system mediated overexpression and knockdown

Tet-On 3G systems are inducible gene expression systems with two elements, namely, the Tet-On 3G transactivator protein and a gene of interest under the control of a TRE3G promoter. The PiggyBac transposon system has been shown to be highly efficient in mediating gene transfer.(Lu & Huang, 2014) Therefore, it has been modified to deliver the multiplex Tet-On 3G system. Briefly, the KANK2 gene sequence and two shRNAs targeting the KANK2-L isoform (shKANK2-L) and KANK2-S isoform (shKANK2-S) were cloned into the pBX-093 plasmid (PB5-HS4- SV40-puro-2A-tetON3G-pA-HS4-TRE-AzaminGreen-2A-Tet3G-RNAi-GpA-HS4- PB3, a kind gift from Dr. Wei Huang, Southern University of Science and Technology, Shenzhen, China) respectively. Cells were subsequently co-transfected with the constructed pBX-093 plasmid and pBX-090 plasmid (pN1- CMV-PGK-piggybac, a kind gift from Dr. Wei Huang) and sorted based on Azamin Green. Before the functional assays, cells were treated with 1μg/ml doxycycline (DOX) to induce the overexpression of KANK2 or knockdown of KANK2-S and KANK2-L. Primers and KANK2-shRNAs sequences are listed in Supplementary file 3.

### RNA extraction, PCR and qRT-PCT analysis

RNA extraction was performed using TRIzol reagent (Vazyme Biotech, Nanjing, China) according to the manufacturer’s instructions. 1μg of total RNA was used to synthesize the first strand of cDNA using Hifair® Ⅲ 1st Strand cDNA Synthesis SuperMix (Yeasen, Shanghai, China, #11141ES60,).

For PCR analysis, Green Taq Mix (#P131-03, Vazyme) was used according to the manufacturer’s instructions. PCR products were analyzed by electrophoresis on a 1% agarose gel.

For qRT-PCR analysis, the SYBR® Green Premix Pro Taq HS qPCR Kit (Accurate Biology, Changsha, China, #AG11701) was used according to the manufacturer’s instructions. The relative changes in gene expression were calculated using the 2 -ΔΔCt method. The primer sequences are listed in Supplementary file 3.

### Cell viability assay, foci formation assay and soft agar colony formation assay

CCK8 assay was performed to analyze cell viability. Cells were digested and re-cultured in 96-well plates at 2000 cells per well in 100μl of medium. Each well was added 10μl CCK8 solution (MCE, New Jersey, USA, HY-K0301) and incubated for 3 h at 37 °C. Optical density was measured using a microplate reader at a wavelength of 452 nm within 5 days. Triplicate repeats were performed to determine the variance and significance. The values were plotted by averaging triplicate wells.

For the foci formation assay, cells were digested and re-cultured in 6-well plates at 500 cells per well in 2 ml medium. After incubation for 9 days, natural monolayer colonies were formed. The cells were subsequently washed with PBS, fixed with 4% paraformaldehyde, stained with purple crystals for 10 min, and washed with PBS three times. The results were photographed, and the number of clones was counted.

For the soft agar colony formation assay, 2000 *PLC-8024* cells were mixed with 10% FBS, 1× DMEM, and 0.35% agar as the upper layer, whereas the bottom layer contained 10% FBS, 1× DMEM and 0.6% agarose. The colonies were photographed under a microscope and the clone numbers were counted 20 days later.

### Cell cycle flow cytometry

Cells were grown to a density of 70% in 6-well dishes and collected for staining with propidium iodide (PI, Sangon, Shanghai, China, #A601112-0100) according to the manufacturer’s instructions. A CytoFLEX cytometer (Beckman Coulter, California, USA) was used to measure changes in cell cycle, and Modfit software was used to analyze the results.

A double thymidine block was used to synchronize cells at the G1/S boundary. Briefly, the cells were treated with 2mM thymidine (Thd, Sigma, #T1895-1G) for 18h and then washed to remove thymidine. After a 9-hour incubation in fresh medium, the cells were treated with the second round of Thd (2 mM) and synchronized at the G1/S boundary. Cells were released by washing with pre-warmed 1x PBS and incubating the cells in pre-warmed fresh media. Cells were collected at 0, 3h for analysis of cell cycle by DNA staining using PI.

To synchronize the cells at the G2/M boundary, the cells were treated with 2mM Thd for 24h and then washed to remove thymidine. After 3-hour incubation in fresh medium, cells were successively treated with 100ng/ml nocodazole (MCE, #HY- 13520) and then synchronized at the G2/M boundary. Cells were released by washing with pre-warmed 1x PBS and incubating the cells in pre-warmed fresh media. Cells were collected at 0, 4h for analysis of cell cycle by DNA staining using PI.

### Biotinylated RNA pulldown and RNA pulldown coupled with mass spectrometry

Biotinylated RNA-pulldown assays were performed as previously described(Panda, Martindale, & Gorospe, 2016). Briefly, exon with 60bp intron sequences at the 5’ or 3’ splice site were cloned into pcDNA3.1, and the plasmid was linearized with the following primers: Forward: TGCTCTGATGCCGCATAGTT, Reverse: GCCCACTACGTGAACCATCA. The linearized DNA fragment was used as a template to generate biotin-labeled RNA for precipitation from *PLC-8024* cell lysate. Subsequently, WB assay was performed to detect precipitation.

RNA pulldown coupled with mass spectrometry was performed as previously described.(Savulescu, Stoychev, Mamputha, & Mhlanga, 2020) Briefly, biotinylated DNA probes were designed to target the exon1-2 junction or the exon1-3 junction of KANK2 and their antisense sequences (exon1-2 sense: CAGCGGCCGGAGCGCGCAAGGTGTTGAAAGACAGAGAAGC, exon1-3 sense: CAGCGGCCGGAGCGCGCAAGGTAAGCCTCAGCCGGTGCTG, exon1-2 antisense: GCTTCTCTGTCTTTCAACACCTTGCGCGCTCCGGCCGCTG, exon1-3 antisense: CAGCACCGGCTGAGGCTTACCTTGCGCGCTCCGGCCGCTG). These probes were incubated with *PLC-8024* cell lysates, followed by an additional incubation with streptavidin beads. After on-bead digestion and desalination, the extracted peptides were analyzed by LC-MSMS.

### RIP

A total of 107 cells were washed with PBS and crosslinked with a 1% formaldehyde solution for 15 min. After neutralization with 0.125M glycine, the cells were lysed in RIPA lysis buffer (Week) (Beyotime Biotechnology, Shanghai, China, #P0013D) for 15 min and ultrasonicated. 0.5μg mouse anti-Flag antibody (Sigma, #F1804) or IgG control (Mouse IgG, Sigma, #I5381-1MG; Rabbit IgG, ThermoFisher, # 10500C) was used for immunoprecipitation. A 10%-volume of lysate was used as the input. RNA in the precipitates and input was subsequently extracted using the acid phenol-chloroform method. The extracted RNAs were detected by qRT-PCR using the Universal Blue qPCR SYBR Green Master Mix (Yeasen, #11184ES08) according to the manufacturer’s instructions. The expression of RIP RNA was calculated as follow(Gagliardi & Matarazzo, 2016): %input(RIP)=2-ΔCt(normalized RIP). ΔCt(normalized RIP)=Average Ct(RIP) - Average Ct(input) - log2(input dilution factor), input dilution factor=(fraction of input RNA)-1. The data were presented as fold enrichment of RNAs in RIP over the IgG control: fold enrichment=% input(RIP)/ % input(IgG)

### RNA seq samples preparation and analysis

USP39 knockdown and control *PLC-8024* cells (three biological replicates per sample) and the tumor or para-tumor tissues collected from USP39HKI and wild-type (WT) mice were suspended in TRIzol reagent (Invitrogen) and sent to Novogene for RNA sequencing. RNA-seq libraries prepared using oligo (dT) beads and rRNA removal methods were pooled and sequenced using an Illumina platform. Paired-end reads were mapped to the Homo sapiens GRCh38(hg38) transcriptome and the Mus musculus reference genome GRCm39(mm39) using the STAR RNA-seq aligner. Alignment files were filtered and sorted using SAMTools. The number of reads mapped to each gene was quantified based on the processed alignment files using htseq-count, and differential expression analysis was performed using DESeq2. Genes with a Padj value < 0.05 and |log2(fold change)|>1 were determined to be differentially expressed. GO and KEGG enrichment analyses were performed using the online tool MSigDB (https://www.gsea-msigdb.org/gsea/msigdb/).

Analysis of pre-RNA splicing efficiency was performed based on the processed alignment files. Briefly, paired-end reads were aligned to the human reference genome GRCh38 using the STAR aligner, and then filtered and sorted using SAMTools. For each splice site, the number of reads covering the first base at the 5’ end of the intron (i.e., 5’ intron coverage), number of reads covering the last base at the 3’ end of the intron (i.e., 3’ intron converage), number of reads covering the last base at the 3’ end of the upstream exon (i.e., 3’ exon coverage), and number of reads covering the first base at the 5’ end of the downstream exon (i.e., 5’ exon coverage) were quantified using bedtools. Splicing efficiencies at the 5’ and 3’ splicing sites were then computed as follows: Efficiency 5’ = 1-(5’ intron coverage)/(3’ exon coverage) and, Efficiency 3’ = 1 – (3’ intron coverage)/(5’ exon coverage). Splice sites with 5’ and 3’ exon coverage lower than 100 were removed from the downstream analysis.

Analysis of alternative splicing and pre-RNA splicing efficiency was performed using rMATS with default parameters. Total counts of reads spanning junctions and reads that did not cross an exon boundary (i.e., outputs in the files [AS_Event]). MATS.JCEC.txt) was used together with the counts of intron reads to identify alternative splicing events. Alternative splicing events with FDR<0.05 and |IncLevelDifference|>0.1 were identified as significant events. Events shared between different experiments were identified based on the genomic coordinates of their associated exons using MATLAB scripts. Genomic coordinates of exons involved in alternative splicing events were retrieved from the rMATS output files using awk.

### Minigene and PCR assay

Upstream, cassette and downstream exons with 60 bp intron sequences at the 5’ and 3’ splice sites were cloned into the pcDNA3.1(-) vector plasmid. Splicing of minigenes was verified by RT-PCR. The primer sequences are listed in Supplementary file 3.

### Statistical analysis

Statistical analyses were performed using SPSS 24.0 or GraphPad Prism 8.0. The mRNA levels of USP39 and KANK2-L isoforms in paired non-tumor and tumor samples were compared using a paired two-tailed Student’s t-test. *USP39* expression levels in unpaired clinical samples were compared using unpaired two-tailed Student’s t-test. *USP39* expression levels in HCC samples with different tumor stages and neoplasm histologic grades were compared using the Kruskal-Wallis test. Differences in overall survival and disease-free survival were calculated using Kaplan-Meier plots and log-rank tests. Correlations between two statistical variables were analyzed using Pearson’s correlation analysis. An unpaired two-tailed Student’s t-test was used to compare observations, such as the number of foci and the relative expression of target genes, between any two preselected groups. Results are expressed as the mean ± standard error of the mean. P<.05 was considered to be statistically significant.

### Code Availability

Code is available on GitHub at https://github.com/VikkiYan/RBP-enrichment.git.

### Data Availability

The data generated in this study are publicly available in Sequence Read Archive (SRA) database at PRJNA855898 (reviewer link: https://dataview.ncbi.nlm.nih.gov/object/PRJNA855898?reviewer=6ulsa5g7sfovp 75ca1dkhufjtt) and PRJNA864041 (reviewer link: https://dataview.ncbi.nlm.nih.gov/object/PRJNA864041?reviewer=bto2vtu69ks5n 94gs8qtlq6guq).The data of mass spectrometry analysis generated in this study are publicly available in the PRIDE database at PXD035928.

## Acknowledgments

This research was supported by the Center for Computational Science and Engineering at Southern University of Science and Technology. We also thank Dr. Wei Huang for generously gifting the pBX093 and pBX090 plasmids.

## Author Contributions

JZ, SW, XO, YL – study concept/design; JZ, SW, MT, PH, LS, YD, HY, FW, HG – experimental data acquisition; YW, JR, HL, ZD– analysis and interpretation of data; SX – provided human samples and clinical data analyses; JZ, SW, ZD, YL – drafted the manuscript; YL–obtained funding, editing, and supervision; all authors have read and edited the manuscript.

---

**Fig. 1-figure supplement 1.**
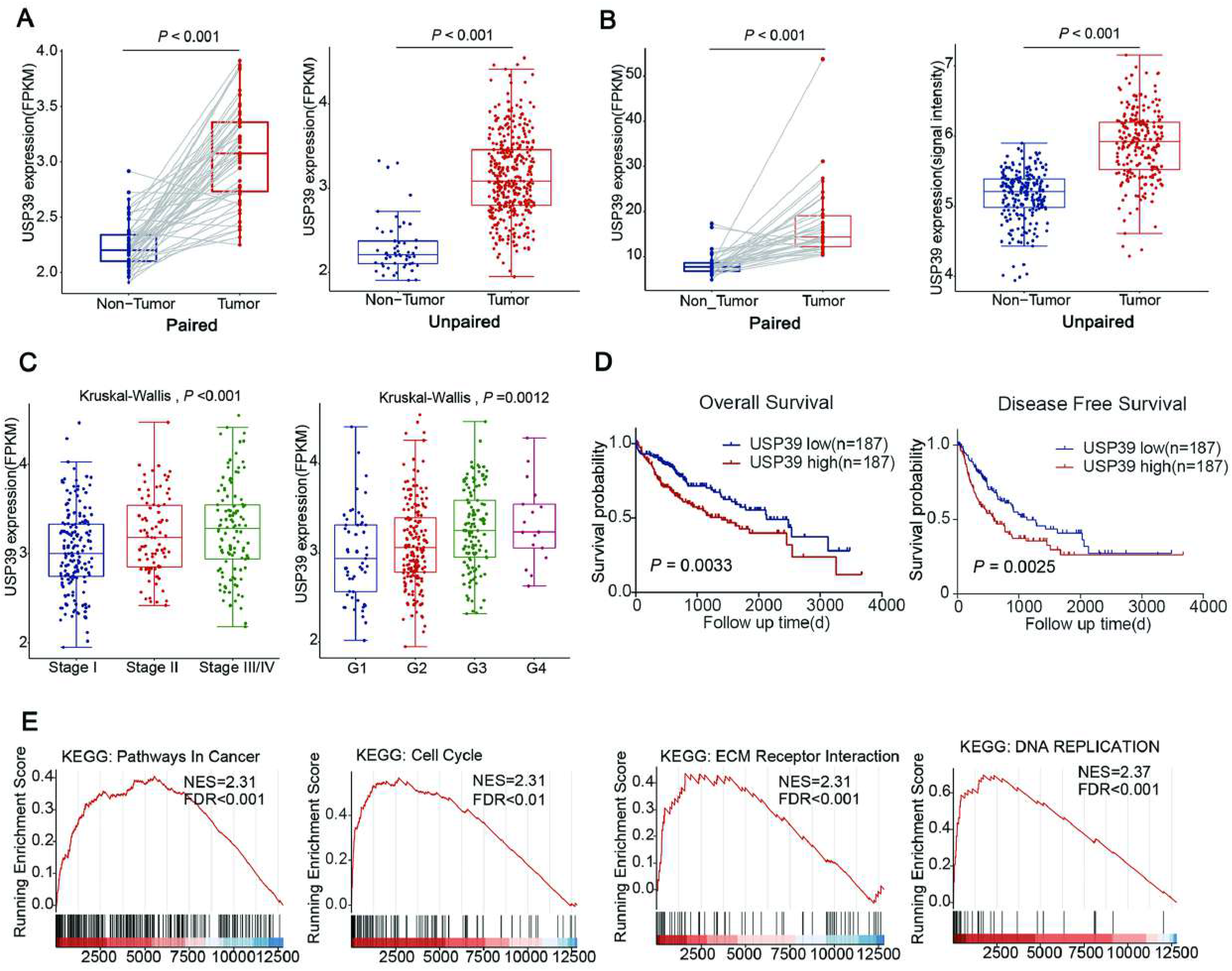
Overexpression of USP39 associates with HCC pathogenesis and aberrant cell cycle signaling. (A) USP39 expression in TCGA-LIHC database, paired and unpaired Student’s t- test. (B) USP39 expression was upregulated in paired (accession no. GSE124535) and unpaired HCC tissues (accession no. GSE14520) in GEO cohorts, paired and unpaired Student’s t test. (C) mRNA levels of USP39 in HCC samples with different tumor stages and neoplasm histologic grades (TCGA-LIHC), Kruskal-Wallis test. (D) Kaplan–Meier OS curves and DFS curves of TCGA HCC patients with high or low USP39 expression (median expression value as a cut-off). (E) GSEA of the indicated gene sets in USP39 high versus low patients from the TCGA-LIHC dataset.

**Fig. 2-figure supplement 1.**
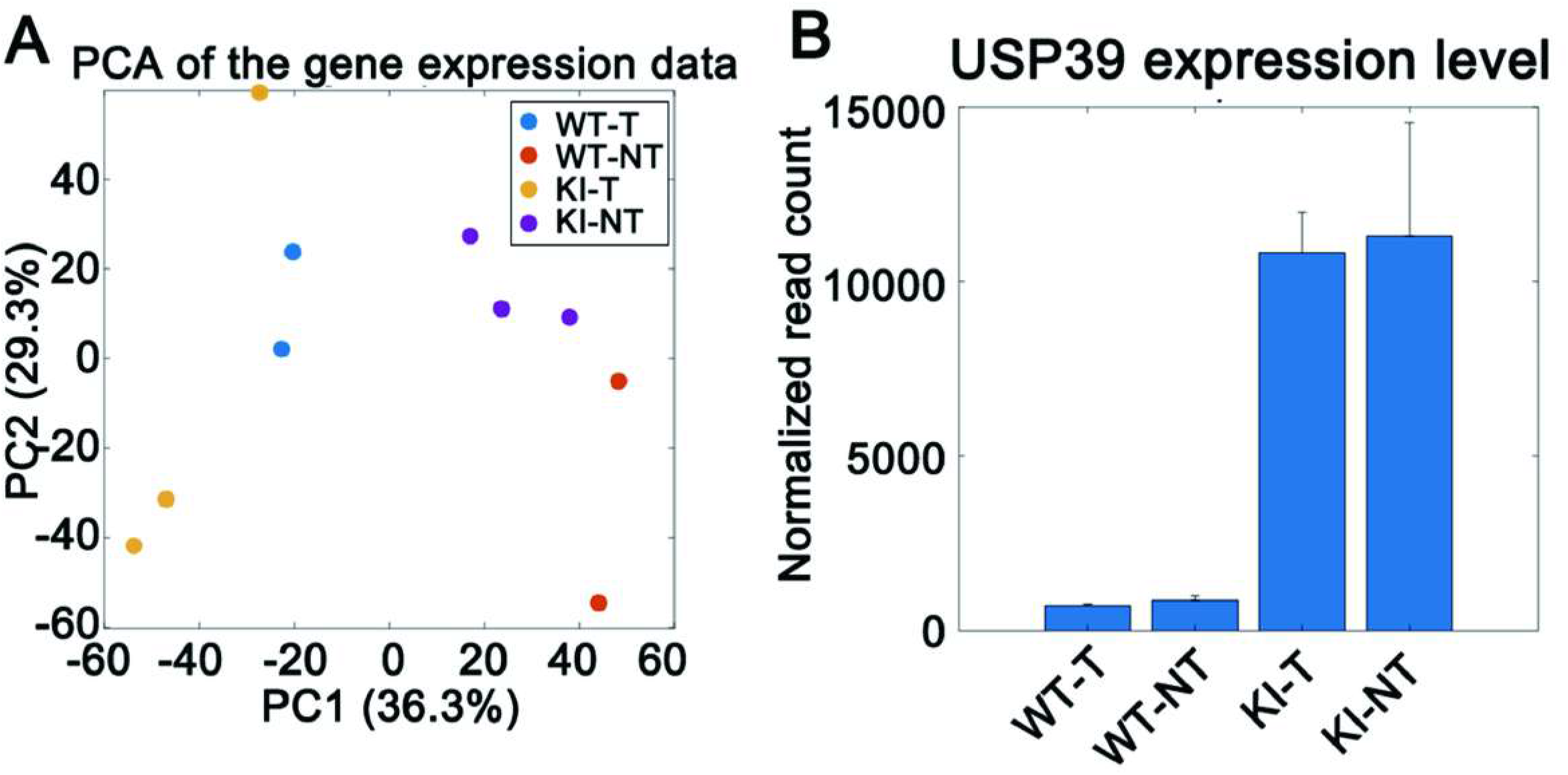
Transcriptome sequencing of hepatocyte- specific USP39 knock-in mice. (A) PCA analysis of wildtype tumors, wildtype non-tumors, knockin tumors and knockin non-tumors. (B) Verification of USP39 overexpression.

**Fig. 3-figure supplement 1.**
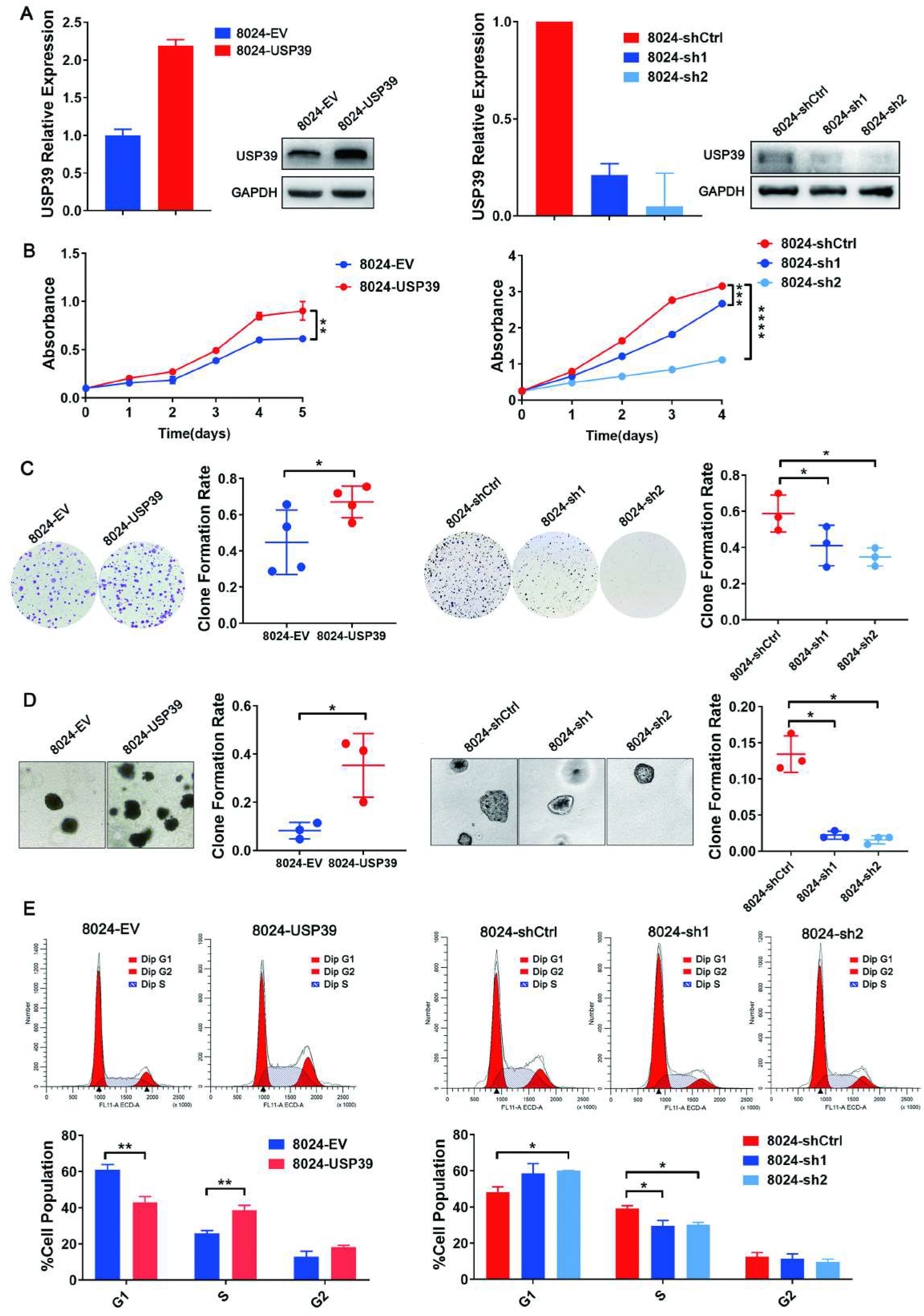
USP39 promotes HCC cell proliferation and cell cycle progression in PLC-8024 cells. (A) Ectopic expression/knockdown of USP39 in PLC-8024 cells was verified by qRT-PCR and WB (EV: Empty Vector, Ctrl: Control). (B) CCK8 assay assessed viability of the indicated cells. (C, D) Representative images and quantification of foci formation (C) or clone formation in soft agar (D) induced by the indicated cells (n=3). (E) Representative flow cytometry histograms of cell cycle progression and statistical results of cell cycle phase distribution (n=3). Mean ± SD. P values were determined using unpaired Student’s t-test. ∗P <.05, ∗ ∗P <.01, ∗∗∗P < .001, ∗∗∗∗P < .0001.

**Fig. 3-figure supplement 2.**
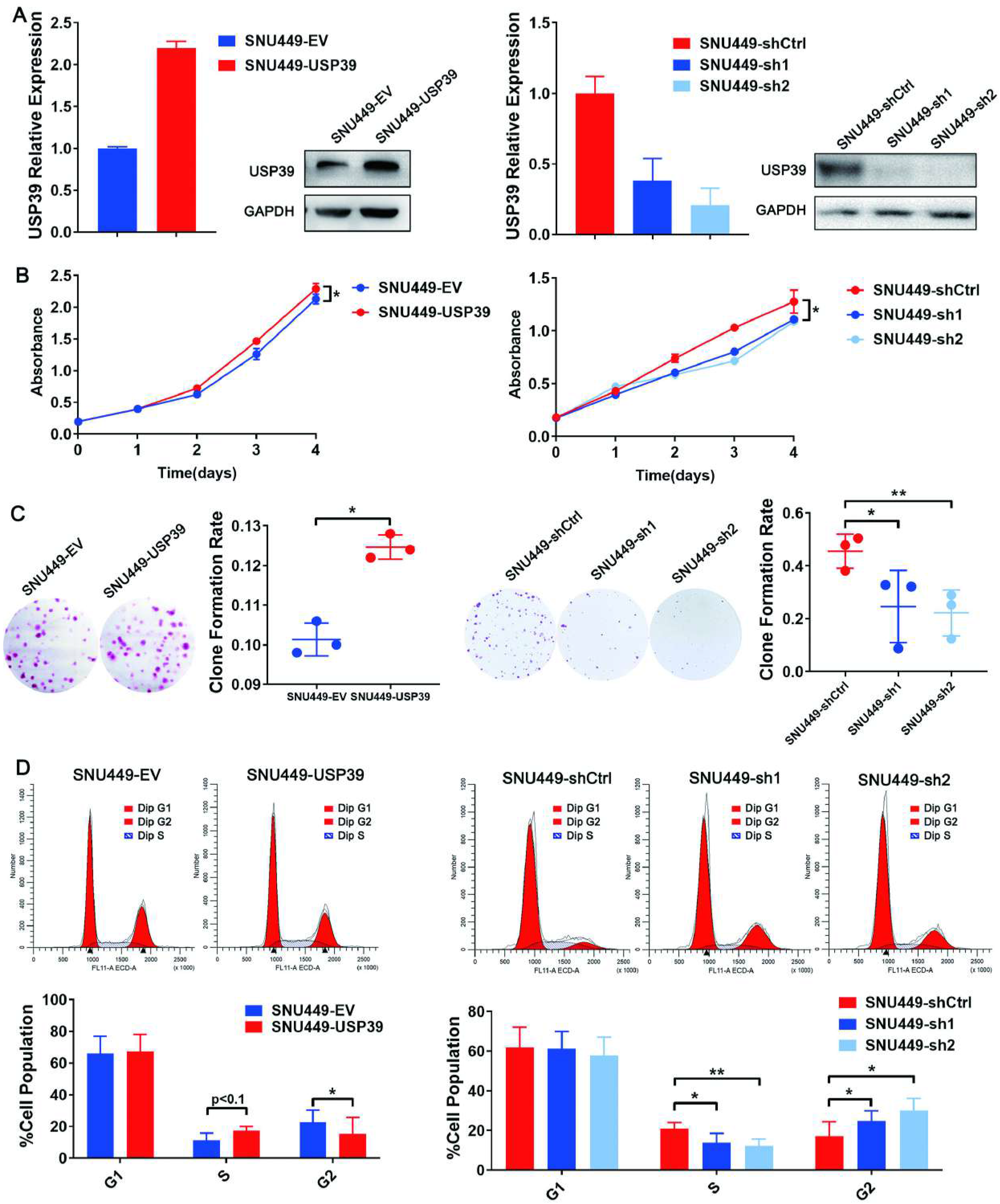
USP39 promotes HCC cell proliferation and cell cycle progression in SNU-449 cells. (A) Ectopic expression/knockdown of USP39 in SNU-449 cells was verified by qRT-PCR and WB (EV: Empty Vector, Ctrl: Control). (B) CCK8 assay revealed that overexpression of USP39 significantly increased cell proliferation, while knockdown of USP39 decreased cell proliferation. (C) Representative images and quantification of foci formation induced by the indicated cells (n=3). (D) Representative flow cytometry histograms of cell cycle progression and statistical results of cell cycle phase distribution (n=3). Mean ± SD. P values by unpaired Student’s t test. ∗P <.05, ∗∗P <.01.

**Fig. 3-figure supplement 3.**
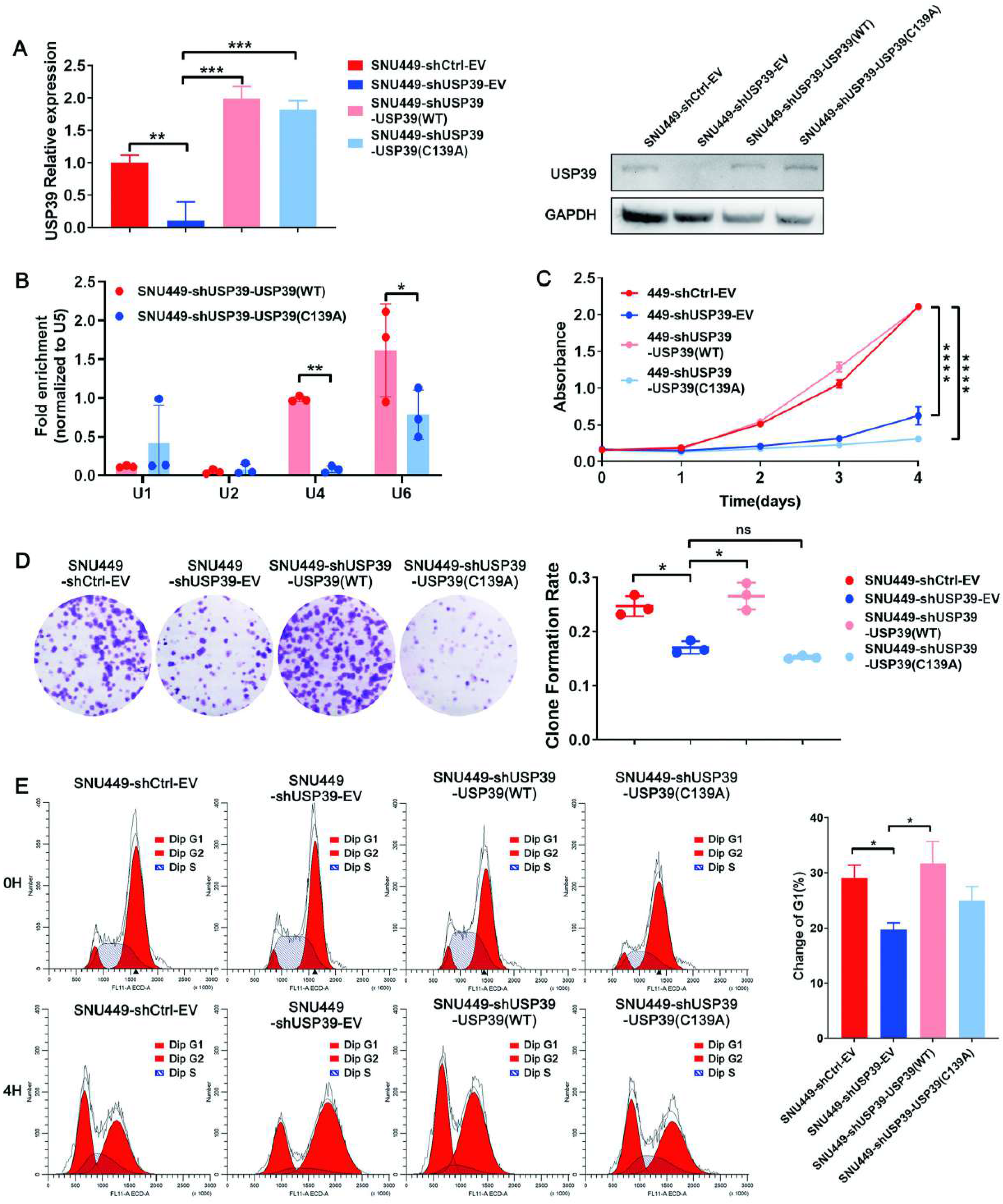
Wild-type USP39 but not C139A mutant rescued the inhibition effect of USP39 deficiency on cell proliferation in SNU-449 cells. (A) Flag-USP39 (WT) and Flag-USP39 (C139A) were introduced to USP39- deficiency (shUSP39) SNU-449 cells and verified with qRT-PCR and WB assays. (B) RIP experiments were performed using antibodies against Flag in the indicated cells. The levels of U1, U2, U4, U6 snRNA in immunoprecipitated RNA were detected using qRT-PCR and normalized to the levels of U5 snRNA in each sample (n=3). (C) CCK8 assay showed that only wild-type USP39, but not C139A mutant, could rescue the inhibition effect of USP39 deficiency on cell proliferation. (D). Representative images and quantification of foci formation induced by the indicated cells (n=3). (E) Cells were synchronized at G2/M boundary after treated with Thd and nocodazole. Following the release, the cells arrested in G2/M (0h) would reenter G1 phase (4h). Cell cycle profiles showed changes in G1 cell populations at 4 hours post-release. C139A mutant failed to rescue cell cycle arrest caused by USP39 knockdown. Mean ± SD. P values by paired Student’s t test (B), or unpaired Student’s t test (A, C-E). ∗P <.05, ∗∗P <.01, ∗∗∗P < .001, ∗∗∗∗P < .0001. ns: not significant.

**Fig. 5-figure supplement 1.**
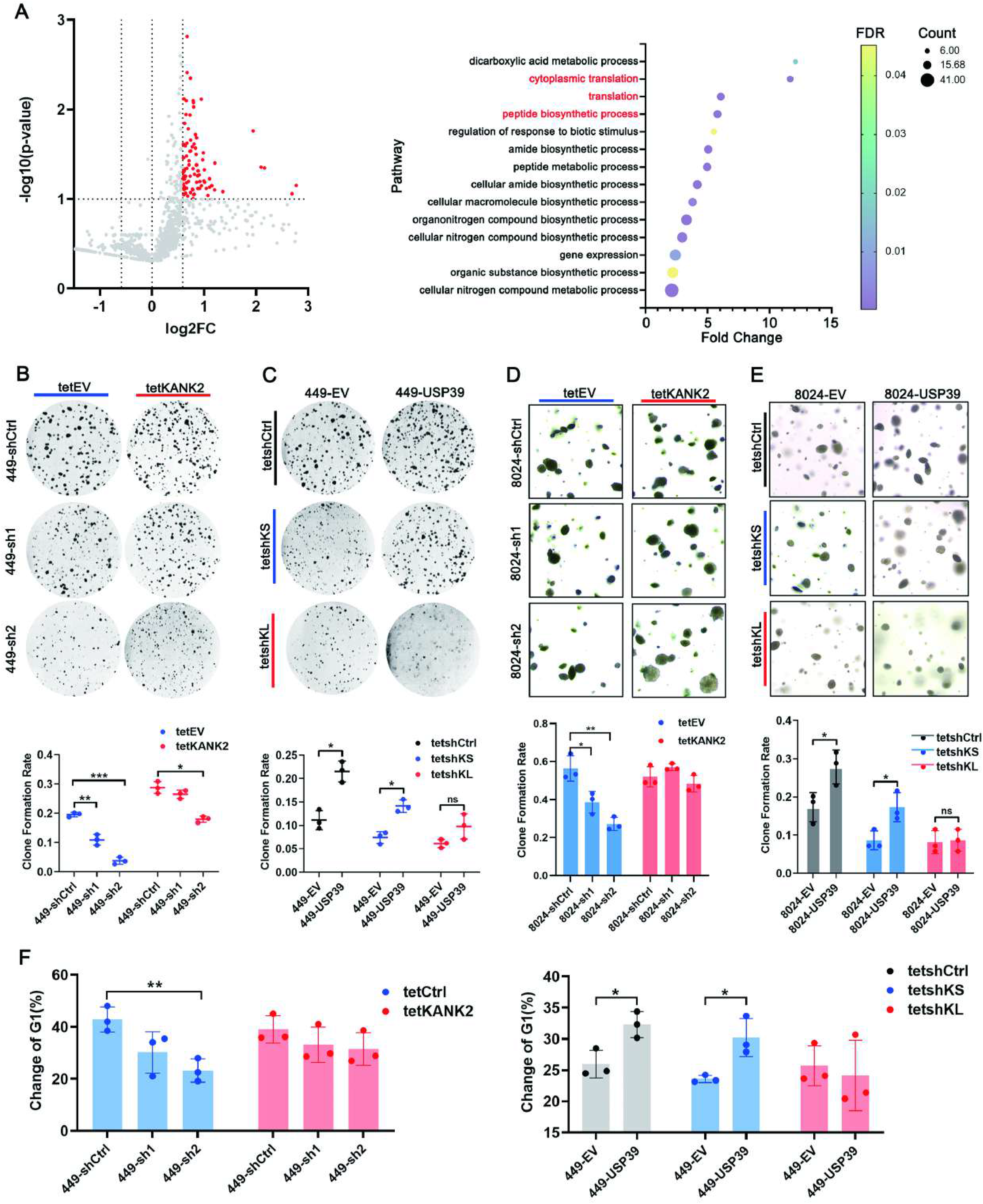
KANK2-L exhibits stronger pro-proliferative activities than KANK2-S. (A) Mass spectrometry analysis of RNA-pulldown samples: red splashes mean significantly enriched genes in KANK2-L pulldown samples (left). The GO BP enrichment analysis of the significantly enriched genes in KANK2-L pulldown samples (right). (B, C) Tet-KANK2 was introduced to USP39-deficient SNU-449 cells (B) and Tet- shKANK2-L(tetshKL) and Tet-KANK2-S (tetshKS) were introduced into USP39- overexpressing SNU-449 cells (C). Cells were treated with 1μg/ml DOX and foci formation assay was performed to access growth of the indicated cells. Representative images and statistical results are shown (n=3). (D, E) Tet-KANK2 was introduced to USP39-deficient PLC-8024 cells (D) and Tet-shKANK2-L(tetshKL) and Tet-KANK2-S (tetshKS) were introduced into USP39-overexpressing PLC-8024 cells (E). After treatment with 1μg/ml DOX, soft agar formation assay was performed to access anchorage-independent growth of the indicated cells. Representative images and statistical results are shown (n=3). (F) Tet-KANK2 was introduced to USP39-deficient SNU-449 cells (left) and Tet- shKANK2-L(tetshKL) and Tet-KANK2-S (tetshKS) were introduced into USP39- overexpressing SNU-449 cells (right). Cells were treated with Thd and nocodazole to synchronize them at G2/M boundary. Following the release, the cells arrested in G2/M (0h) would reenter G1 phase (4h). Cell cycle profiles showed changes in G1 cell populations at 4 hours post-release (n=3). Mean ± SD. p values by Pearson’s correlation coefficient (A) and unpaired Student’s t test (B-F). ∗P < .05, ∗∗P < .01, ∗∗∗P < .001. ns: not significant.

**Fig. 5-figure supplement 2.**
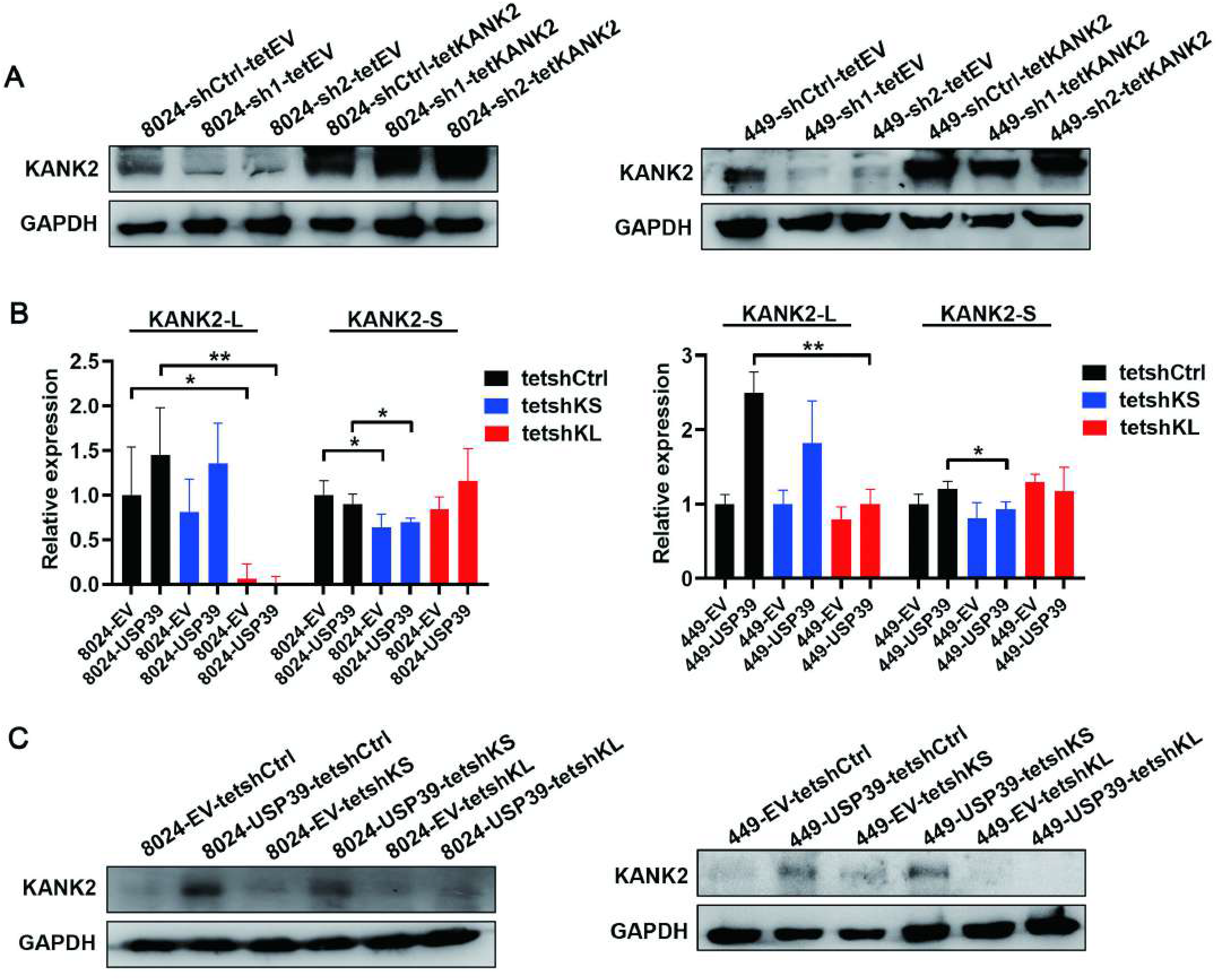
Tet-On system mediates KANK2 overexpression and isoform-specific knockdown. (A) Tet-On system mediated KANK2 overexpression in USP39-deficient PLC- 8024 and SNU-449 cells. The overexpression effects were verified by WB. (B, C) Tet-shKANK2-L(tetshKL) and Tet-KANK2-S (tetshKS) were introduced into USP39-overexpressing PLC-8024 and SNU-449 cells. The knockdown effects were examined by qRT-PCR (B) and WB (C). KANK2-L depletion reduced KANK2 protein expression, while KANK2-S targeting barely affected KANK2 protein expression despite successful knockdown of KANK2-S mRNA.

**Supplementary file 1: RBP motif enrichment analyses of 5’ and 3’ splice junction sequences (cassette exons vs backgroup exons). Related to** **Figure 6**.

**Supplementary file 2: Demographic information of the in-house HCC cohort.**

**Supplementary file 3: The sequences of PCR primers, qPCR primers, siRNA and shRNA.**

## References

Alsafadi, S., Houy, A., Battistella, A., Popova, T., Wassef, M., Henry, E., . . . Stern, M.- H. (2016). Cancer-associated SF3B1 mutations affect alternative splicing by promoting alternative branchpoint usage. Nat Commun, 7(1), 10615. doi:10.1038/ncomms10615

Barash, Y., Calarco, J. A., Gao, W., Pan, Q., Wang, X., Shai, O., . . . Frey, B. J. (2010). Deciphering the splicing code. Nature, 465(7294), 53–59. doi:10.1038/nature09000

Bertram, K., Agafonov, D. E., Dybkov, O., Haselbach, D., Leelaram, M. N., Will, C. L., . . . Stark, H. (2017). Cryo-EM Structure of a Pre-catalytic Human Spliceosome Primed for Activation. Cell, 170(4), 701–713.e711. doi:10.1016/j.cell.2017.07.011

Chen, H., Gao, F., He, M., Ding, X. F., Wong, A. M., Sze, S. C., . . . Wong, N. (2019). Long-Read RNA Sequencing Identifies Alternative Splice Variants in Hepatocellular Carcinoma and Tumor-Specific Isoforms. Hepatology, 70(3), 1011–1025. doi:10.1002/hep.30500

Chen, J., & Weiss, W. A. (2015). Alternative splicing in cancer: implications for biology and therapy. Oncogene, 34(1), 1–14. doi:10.1038/onc.2013.570

Danan-Gotthold, M., Golan-Gerstl, R., Eisenberg, E., Meir, K., Karni, R., & Levanon, E. Y. (2015). Identification of recurrent regulated alternative splicing events across human solid tumors. Nucleic Acids Res, 43(10), 5130–5144. doi:10.1093/nar/gkv210

Ding, K., Ji, J., Zhang, X., Huang, B., Chen, A., Zhang, D., . . . Wang, J. (2019). RNA splicing factor USP39 promotes glioma progression by inducing TAZ mRNA maturation. Oncogene, 38(37), 6414–6428. doi:10.1038/s41388-019-0888-1

Dong, L., Yu, L., Li, H., Shi, L., Luo, Z., Zhao, H., . . . Lin, Z. (2020). An NAD(+)-Dependent Deacetylase SIRT7 Promotes HCC Development Through Deacetylation of USP39. iScience, 23(8), 101351. doi:10.1016/j.isci.2020.101351

Dong, X., Liu, Z., Zhang, E., Zhang, P., Wang, Y., Hang, J., & Li, Q. (2021). USP39 promotes tumorigenesis by stabilizing and deubiquitinating SP1 protein in hepatocellular carcinoma. Cell Signal, 85, 110068. doi:10.1016/j.cellsig.2021.110068

Eskens, F. A., Ramos, F. J., Burger, H., O’Brien, J. P., Piera, A., de Jonge, M. J., . . . Tabernero, J. (2013). Phase I pharmacokinetic and pharmacodynamic study of the first-in-class spliceosome inhibitor E7107 in patients with advanced solid tumors. Clin Cancer Res, 19(22), 6296–6304. doi:10.1158/1078-0432.Ccr-13-0485

Eymin, B. (2021). Targeting the spliceosome machinery: A new therapeutic axis in cancer? Biochem Pharmacol, 189, 114039. doi:10.1016/j.bcp.2020.114039

Fraile, J. M., Manchado, E., Lujambio, A., Quesada, V., Campos-Iglesias, D., Webb, T. R., . . . Freije, J. M. (2017). USP39 Deubiquitinase Is Essential for KRAS Oncogene-driven Cancer. J Biol Chem, 292(10), 4164–4175. doi:10.1074/jbc.M116.762757

Fu, X. D., & Ares, M., Jr. (2014). Context-dependent control of alternative splicing by RNA-binding proteins. Nat Rev Genet, 15(10), 689–701. doi:10.1038/nrg3778

Gagliardi, M., & Matarazzo, M. R. (2016). RIP: RNA Immunoprecipitation. Methods Mol Biol, 1480, 73–86. doi:10.1007/978-1-4939-6380-5_7

Gao, Q., Zhu, H., Dong, L., Shi, W., Chen, R., Song, Z., . . . Fan, J. (2019). Integrated Proteogenomic Characterization of HBV-Related Hepatocellular Carcinoma. Cell, 179(2), 561–577.e522. doi:10.1016/j.cell.2019.08.052

Gee, H. Y., Zhang, F., Ashraf, S., Kohl, S., Sadowski, C. E., Vega-Warner, V., . . . Hildebrandt, F. (2015). KANK deficiency leads to podocyte dysfunction and nephrotic syndrome. J Clin Invest, 125(6), 2375–2384. doi:10.1172/jci79504

Geuens, T., Bouhy, D., & Timmerman, V. (2016). The hnRNP family: insights into their role in health and disease. Hum Genet, 135(8), 851–867. doi:10.1007/s00439-016-1683-5

Hadjivassiliou, H., Rosenberg, O. S., & Guthrie, C. (2014). The crystal structure of S. cerevisiae Sad1, a catalytically inactive deubiquitinase that is broadly required for pre-mRNA splicing. Rna, 20(5), 656–669. doi:10.1261/rna.042838.113

Huang, Y., Pan, X. W., Li, L., Chen, L., Liu, X., Lu, J. L., . . . Cui, X. G. (2016). Overexpression of USP39 predicts poor prognosis and promotes tumorigenesis of prostate cancer via promoting EGFR mRNA maturation and transcription elongation. Oncotarget, 7(16), 22016–22030. doi:10.18632/oncotarget.7882

Huang, Y. H., Chung, C. S., Kao, D. I., Kao, T. C., & Cheng, S. C. (2014). Sad1 counteracts Brr2-mediated dissociation of U4/U6.U5 in tri-snRNP homeostasis. Mol Cell Biol, 34(2), 210–220. doi:10.1128/mcb.00837-13

Li, X., Yuan, J., Song, C., Lei, Y., Xu, J., Zhang, G., . . . Song, G. (2021). Deubiquitinase USP39 and E3 ligase TRIM26 balance the level of ZEB1 ubiquitination and thereby determine the progression of hepatocellular carcinoma. Cell Death Differ, 28(8), 2315–2332. doi:10.1038/s41418-021-00754-7

Liao, Y., Li, L., Liu, H., & Song, Y. (2021). High Expression of Ubiquitin-Specific Protease 39 and Its Roles in Prognosis in Patients with Hepatocellular Carcinoma. Evid Based Complement Alternat Med, 2021, 6233175. doi:10.1155/2021/6233175

Lu, X., & Huang, W. (2014). PiggyBac mediated multiplex gene transfer in mouse embryonic stem cell. PLoS One, 9(12), e115072. doi:10.1371/journal.pone.0115072

Lygerou, Z., Christophides, G., & Séraphin, B. (1999). A novel genetic screen for snRNP assembly factors in yeast identifies a conserved protein, Sad1p, also required for pre-mRNA splicing. Mol Cell Biol, 19(3), 2008–2020. doi:10.1128/mcb.19.3.2008

Ni, W., Bian, S., Zhu, M., Song, Q., Zhang, J., Xiao, M., & Zheng, W. (2021). Identification and Validation of Ubiquitin-Specific Proteases as a Novel Prognostic Signature for Hepatocellular Carcinoma. Front Oncol, 11, 629327. doi:10.3389/fonc.2021.629327

Pan, Z., Pan, H., Zhang, J., Yang, Y., Liu, H., Yang, Y., . . . Zhou, W. (2015). Lentivirus mediated silencing of ubiquitin specific peptidase 39 inhibits cell proliferation of human hepatocellular carcinoma cells in vitro. Biol Res, 48(1), 18. doi:10.1186/s40659-015-0006-y

Panda, A. C., Martindale, J. L., & Gorospe, M. (2016). Affinity Pulldown of Biotinylated RNA for Detection of Protein-RNA Complexes. Bio-protocol, 6(24), e2062. doi:10.21769/BioProtoc.2062

Papasaikas, P., Tejedor, J. R., Vigevani, L., & Valcárcel, J. (2015). Functional splicing network reveals extensive regulatory potential of the core spliceosomal machinery. Mol Cell, 57(1), 7–22. doi:10.1016/j.molcel.2014.10.030

Pei, Q., Ni, W., Yuan, Y., Yuan, J., Zhang, X., & Yao, M. (2022). HSP70 Ameliorates Septic Lung Injury via Inhibition of Apoptosis by Interacting with KANK2. Biomolecules, 12(3). doi:10.3390/biom12030410

Savulescu, A. F., Stoychev, S., Mamputha, S., & Mhlanga, M. M. (2020). Biochemical Pulldown of mRNAs and Long Noncoding RNAs from Cellular Lysates Coupled with Mass Spectrometry to Identify Protein Binding Partners. Bio-protocol, 10(11), e3639. doi:10.21769/BioProtoc.3639

Seiler, M., Peng, S., Agrawal, A. A., Palacino, J., Teng, T., Zhu, P., . . . Yu, L. (2018). Somatic Mutational Landscape of Splicing Factor Genes and Their Functional Consequences across 33 Cancer Types. Cell Rep, 23(1), 282–296.e284. doi:10.1016/j.celrep.2018.01.088

Seiler, M., Yoshimi, A., Darman, R., Chan, B., Keaney, G., Thomas, M., . . . Buonamici, S. (2018). H3B-8800, an orally available small-molecule splicing modulator, induces lethality in spliceosome-mutant cancers. Nat Med, 24(4), 497–504. doi:10.1038/nm.4493

Shepard, P. J., & Hertel, K. J. (2009). The SR protein family. Genome Biol, 10(10), 242. doi:10.1186/gb-2009-10-10-242

Wang, D., Liang, J., Zhang, Y., Gui, B., Wang, F., Yi, X., . . . Shang, Y. (2012). Steroid receptor coactivator-interacting protein (SIP) inhibits caspase-independent apoptosis by preventing apoptosis-inducing factor (AIF) from being released from mitochondria. J Biol Chem, 287(16), 12612–12621. doi:10.1074/jbc.M111.334151

Wang, S., Wang, Z., Li, J., Qin, J., Song, J., Li, Y., . . . Liu, Z. (2021). Splicing factor USP39 promotes ovarian cancer malignancy through maintaining efficient splicing of oncogenic HMGA2. Cell Death Dis, 12(4), 294. doi:10.1038/s41419-021-03581-3

Will, C. L., & Lührmann, R. (2011). Spliceosome structure and function. Cold Spring Harb Perspect Biol, 3(7). doi:10.1101/cshperspect.a003707

Xiao, Y., Ma, W., Hu, W., Di, Q., Zhao, X., Ma, X., . . . Chen, W. (2022). Ubiquitin-specific peptidase 39 promotes human glioma cells migration and invasion by facilitating ADAM9 mRNA maturation. Mol Oncol, 16(2), 388–404. doi:10.1002/1878-0261.12958

Yu, L., Kim, J., Jiang, L., Feng, B., Ying, Y., Ji, K.-y., . . . Xu, Y. (2020). MTR4 drives liver tumorigenesis by promoting cancer metabolic switch through alternative splicing. Nat Commun, 11(1), 708. doi:10.1038/s41467-020-14437-3

Yuan, X., Sun, X., Shi, X., Jiang, C., Yu, D., Zhang, W., . . . Ding, Y. (2015). USP39 promotes the growth of human hepatocellular carcinoma in vitro and in vivo. Oncol Rep, 34(2), 823–832. doi:10.3892/or.2015.4065

Zhao, Y., Geng, H., Liu, G., Ji, Q., Cheng, X., Li, X., . . . Liu, X. (2021). The Deubiquitinase USP39 Promotes ESCC Tumorigenesis Through Pre-mRNA Splicing of the mTORC2 Component Rictor. Front Oncol, 11, 667495. doi:10.3389/fonc.2021.667495

Zheng, X., Peng, Q., Wang, L., Zhang, X., Huang, L., Wang, J., & Qin, Z. (2020). Serine/arginine-rich splicing factors: the bridge linking alternative splicing and cancer. Int J Biol Sci, 16(13), 2442–2453. doi:10.7150/ijbs.46751

Zhou, X., Wang, R., Li, X., Yu, L., Hua, D., Sun, C., . . . Yu, S. (2019). Splicing factor SRSF1 promotes gliomagenesis via oncogenic splice-switching of MYO1B. J Clin Invest, 129(2), 676–693. doi:10.1172/jci120279

Zhu, X., Ma, J., Lu, M., Liu, Z., Sun, Y., & Chen, H. (2022). The Deubiquitinase USP39 Promotes Esophageal Squamous Cell Carcinoma Malignancy as a Splicing Factor. Genes (Basel), 13(5). doi:10.3390/genes13050819

